# Light environment and seasonal variation in the visual system of the red shiner (*Cyprinella lutrensis*)

**DOI:** 10.1101/2024.05.02.592238

**Authors:** Tarah N. Foster, Alyssa G. Williamson, Bradley R. Foster, Matthew B. Toomey

**Affiliations:** Department of Biological Sciences, University of Tulsa, Oklahoma, 74104

## Abstract

The light environment underwater can vary dramatically over space and time, challenging the visual systems of aquatic organisms. To meet these challenges, many species shift their spectral sensitivities through changes in visual pigment chromophore and opsin expression. The red shiner (*Cyprinella lutrensis*) is a cyprinid minnow species that has rapidly expanded its range throughout North America and inhabits a wide range of aquatic habitats. We hypothesized that visual system plasticity has contributed to the red shiner’s success. We investigated plasticity in chromophore usage and opsin expression by collecting red shiners from three Oklahoma creeks that vary in turbidity throughout the year. We characterized the light environment by spectroradiometry, measured chromophore composition of the eyes with high performance liquid chromatography, characterized CYP27C1 enzyme function through heterologous expression, and examined ocular gene expression by RNA sequencing and *de novo* transcriptome assembly. We observed significantly higher proportions of the long-wavelength shifted A_2_ chromophore in the eyes of fish from the turbid site and in samples collected in winter, suggesting that there may be a temperature-dependent trade-off between chromophore-based spectral tuning and chromophore-related noise. Opsin expression varied between turbid and clear creeks, but did not align with light environment as expected, and the magnitude of these differences was limited compared to the differences in chromophore composition. We confirmed that red shiner *CYP27C1* catalyzes the conversion of A_1_ to A_2_, but the ocular expression of *CYP27C1* was not well correlated with A_2_ levels in the eye, suggesting conversion may be occurring outside of the eye.

## Introduction

Many aquatic animals rely on vision, yet these organisms may face great variation in their visual environment. The spectrum and intensity of light in the aquatic environment varies with depth and is altered by the presence of sediments and dissolved organic material (Cronin et al., 2014; Lythgoe, 1980; Lythgoe, 1988). There is a wealth of evidence indicating that the spectral sensitivities of aquatic organisms have evolved to match their specific light environment (Bowmaker et al., 1994; Margetts et al., 2024) and the ‘sensitivity hypothesis’ predicts that this matching facilitates vision (Lythgoe, 1980; Marshall et al., 2015). The strength of this relationship has made the visual systems of fish an important model system for investigations linking molecular change to ecological and evolutionary processes (Carleton et al., 2005; Fuller et al., 2005; Rennison et al., 2016; Seehausen et al., 2008).

The aquatic light environment is dynamic with predictable (e.g. seasonal shifts in productivity) and unpredictable (e.g. precipitation and turbidity) shifts in light intensity and spectrum. Matching this dynamic light environment presents a special challenge requiring plasticity within the visual system. The spectral sensitivities of fish are primarily determined by the visual pigments of the photoreceptors that consist of the G-protein-coupled receptor, opsin, and the vitamin A-derived molecule bound to it, the chromophore (Carleton et al., 2020; Corbo, 2021). Spectral sensitivity can be fine-tuned over the lifetime of an individual through: 1) changes in the type and level of opsin expressed within a photoreceptor, and/or 2) changes to the chromophore component of the visual pigment. Opsin expression has been extensively studied in multiple model organisms demonstrating both environmentally induced plasticity and heritable genetic determinants (Carleton, 2009; Fuller and Claricoates, 2011; Fuller et al., 2005; Härer et al., 2017; Hofmann et al., 2010; Karagic et al., 2018; Nandamuri et al., 2017; Rennison et al., 2016; Veen et al., 2017). In contrast, the drivers and consequences of chromophore plasticity are less well understood.

The chromophore component of the visual pigment absorbs photons of light and isomerizes, causing a conformational change of the opsin that initiates the phototransduction cascade, ultimately leading to light perception. Many freshwater fish species use two different chromophore molecules, vitamin A_1_-derived 11-*cis* retinal (A_1_) and vitamin A_2_-derived 11-*cis* 3,4 didehydroretinal (A_2_). The difference between the vitamin A_1_ and vitamin A_2_ chromophores is an additional double bond within the terminal β-ionone ring of A_2_ (Corbo, 2021). When combined with many opsins, the additional double bond in A_2_ results in a long-wavelength shift in the sensitivity of the visual pigment, relative to an A_1_-containing pigment. The largest A_1_ to A_2_ shifts in sensitivity are observed among longer-wavelength sensitive pigments (Corbo, 2021; Dartnall and Lythgoe, 1965; Liebman and Entine, 1968; Munz and Schwanzara, 1967). Studies in zebrafish (*Danio rerio*), have revealed that the enzyme cytochrome P450 27c1 (CYP27C1) mediates the conversion A_1_ to A_2_ within the retinal pigment epithelium of the eye (Enright et al., 2015). CYP27C1-mediated chromophore switching is present in sea lamprey (*Petromyzon marinus*) and likely widely conserved among vertebrates (Morshedian et al., 2017).

Chromophore switching appears to be a widely expressed mechanism of visual system plasticity, with shifts observed as animals migrate between different light environments or as the environment changes with the seasons (Bridges, 1972; Corbo, 2021). For example, anadromous salmonids increase levels of vitamin A_2_ chromophore as adults migrate from the blue waters of the open ocean to the long-wavelength shifted environments of their freshwater spawning sites (Alexander et al., 1994; Beatty, 1966). Similarly, catadromous sea lamprey shift from A_2_ to an A_1_ chromophore as they migrate from their natal freshwater streams and rivers to the relatively blue-shifted waters of ocean spawning grounds (Crescitelli, 1956; Morshedian et al., 2017; Wald, 1957).

Among non-migratory freshwater fish species, increased levels of vitamin A_2_ chromophore are associated with increased water turbidity (Bridges, 1964; Bridges, 1965; Bridges, 1972; Escobar-Camacho et al., 2019). These shifts in chromophore composition are thought to be an adaptive plastic response that allow the animals to match new or changed light environment.

The adaptive significance of seasonal shifts in chromophore composition are more difficult to discern. Many freshwater fish species increase the A_2_ chromophore content of their eyes in the colder winter months, compared to the warmer summer months (Bridges, 1964; Corbo, 2021; Temple et al., 2006; Whitmore and Bowmaker, 1989). It has been hypothesized that these seasonal shifts track a red-shift in the aquatic light environments resulting from the positioning of the sun and abundance of suspended particles (Bridges, 1972). However, there is less primary productivity in winter and there may be less spectral filtering by plankton and algae which may result in a relatively blue-shifted light environment. Therefore, it is difficult to predict how light environment will change seasonally and it is likely to vary from site to site. Seasonal changes in temperature may have a direct impact on chromophore function. Visual pigments with the A_2_ chromophore are less thermally stable than the A_1_-containing visual pigments (Ala-Laurila et al., 2007; Donner, 2020). Therefore, A_2_-containing visual pigments are more likely to isomerize and activate the phototransduction cascade in the absence of light. Thus, it may be disadvantageous to utilize A_2_ in warmer waters when the probability of thermal isomerization is high. In cold waters, the probability of thermal isomerization is lower, which may reduce the inherent noise associated with A_2_ chromophore. The extent to which red-shifted longer wavelength light environments versus colder water temperatures induce shifts to a vitamin A_2_-derived chromophore remains unresolved. Thus, chromophore composition plasticity may reflect a trade-off between signal-to-noise ratio of the photoreceptor response and spectral sensitivity that is dependent on both light environment and temperature.

In this study, we examined the associations between visual pigment chromophore composition, light environment, and season in the red shiner (*Cyprinella lutrensis*), a cyprinid minnow species native to the Mississippi river basin of North America (Nico et al. 2019). The red shiner is an invasive species that has rapidly expanded its range throughout North America (Mapping the potential distribution of the invasive Red Shiner, Cyprinella lutrensis (Teleostei: Cyprinidae) across waterways of the conterminous United States). Red shiners thrive in waters varying widely in available light spectra, with mean of yearly median habitat turbidity measures ranging from 10 to 140 nephelometric turbidity units (NTU) and withstand temperatures ranging from 0°C-37°C (Dekar et al., 2014; Dugas and Franssen, 2012; Matthews, 1986; Matthews and Hill, 1977). Visual signaling is an important part of the biology of red shiners, with males displaying elaborate nuptial coloration (Dugas and Franssen, 2011).

This nuptial coloration is more intensely displayed in more turbid habitats (Dugas and Franssen, 2011). The success of invasive red shiners may reflect their capacity for phenotypical plasticity including in the visual system. Red shiners sampled from turbid waters have relatively larger eyes, and fish reared in turbid conditions express higher proportions of long-wavelength sensitive opsin (*LWS)* in their retinas (Chang and Yan, 2019; Dugas and Franssen, 2012). However, it is not known if red shiners also switch chromophore composition or how this switch is influenced by the environmental conditions this species encounters throughout the seasons. To address this question, we sampled red shiners throughout the year from creeks with differing light environments. We used high performance liquid chromatography to characterize the chromophore composition of the retina and RNA sequencing to determine opsin expression. We also characterized the enzymatic function of *CYP27C1* from the red shiner utilizing a heterologous expression system.

## Materials and Methods

### Field Sampling

We sampled red shiners from Big Creek (36.785775, -95.469693), Lightning Creek (36.655701, -95.464655), and Polecat Creek (36.017335, -95.985103) in northeastern Oklahoma, USA from July 2020 to May 2022 (Fig. S1). These sites are part of the Oklahoma Water Resources Board (OWRB) long-term monitoring efforts and turbidity measures have been collected over multiple years. These measures indicate that there are significant differences in turbidity among the sites and across the seasons (site: F_2,77_ = 5.60, P = 0.0054, season: F_3,77_ = 3.22, P = 0.027, Fig. 1a, Fig. S1). Polecat

**Figure 1.**
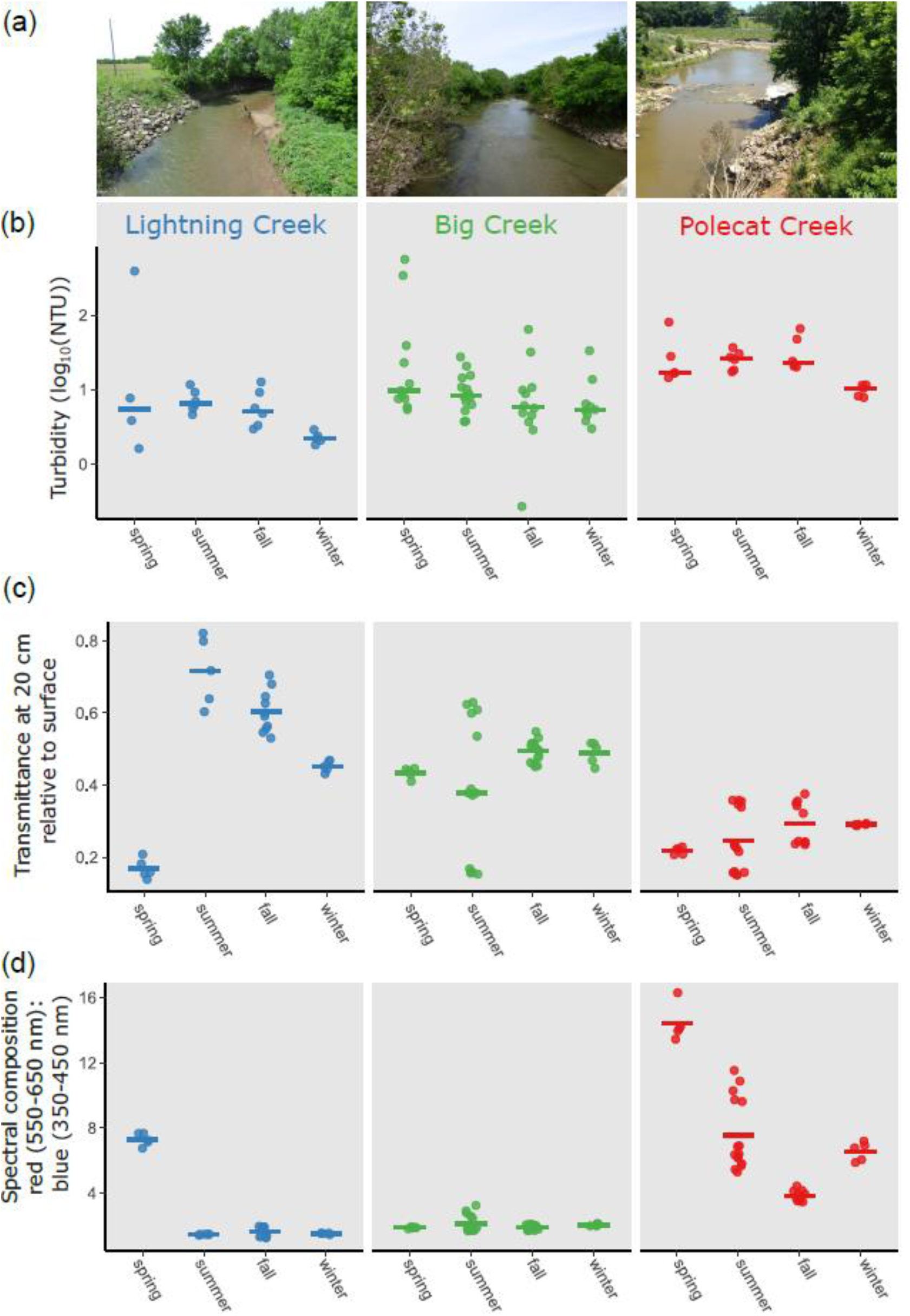
The light environments of our study sites differ in turbidity, transmittance, and spectral composition. (a) Representative images of our study sites: Lighting Creek (left), Big Creek (middle), and Polecat Creek (right, photo: Meridian Engineering). (b) Turbidity of each site measured in Nephelometric Turbidity Units (NTU) for the years spanning from 2011-2018. The bar indicates the mean turbidity in each site and season. Turbidity data provided courtesy of the Oklahoma Conservation Commission. (c) The transmittance measured as the total irradiance 350-700 nm wavelength at 20 cm depth divided by total irradiance at the water’s surface. The bar indicates the mean irradiance at each site and season. (d) The spectral composition of light at 20 cm depth measured as the ratio of red irradiance (550-650 nm) to blue irradiance (350-450 nm). The bar indicates the mean red:blue ratio at each site in each season.

Creek was significantly more turbid than both Big Creek and Lightning Creek (Tukey HSD, P < 0.012) and turbidity is significantly greater in spring compared to the winter season (Tukey HSD, P = 0.0032). Fish were caught via minnow trap, dip net, or by seining in accordance with method of take regulations set by the Oklahoma Department of Wildlife Conservation under a scientific collector’s permit (#10249373). Each site was sampled 7 times during the study, and we collected a minimum of four and a maximum of 28 individuals in each sampling bout. Fish were either euthanized on site or kept alive in an aerated minnow bucket for no more than 1 hour and transported to the University of Tulsa for subsequent euthanasia via overdose of Tricaine-S (MS222) and set on ice (in either case). We immediately enucleated right and left eyes and transferred them to a cooler containing dry ice (in the case of field dissection) or a -80° C freezer (in the case of lab dissection). All samples were stored at -80° C until further analysis. All methods were approved by the University of Tulsa Institutional Animal Care and Use Committee (protocol number TU-0052). Eye diameter and standard body length were measured post-mortem, to the nearest mm, with digital calipers. Fish sampled during breeding season were sexed based on nuptial coloration and via dissection and observation of testes or ovaries (Dugas and Franssen, 2011). Outside of the breeding season, it was not possible to definitively determine sex from morphology.

### Light Measurements

When possible, we collected downwelling irradiance measures at the water’s surface and at 20cm depth during each fish sampling bout. We used an Ocean Optics Flame-S (serial #FLMS14357) miniature spectrometer coupled to an Ocean Insight CC-3-UV-S- Cosine Corrector with Spectralon diffusing material for irradiance measures. We calibrated the spectrophotometer with a radiometric calibrated light source (HL-3P-CAL, Ocean optics Inc.) and initial measures were recorded in units of irradiance (μWatt cm_-2_). These data were processed in R (Team, 2021) utilizing packages Pavo (Maia et al., 2019) and Tidyverse (Wickham et al., 2019). Irradiance measures were converted to units of photon flux (μmol s_-1_ m_-2_) and then summed from 300-700 nm to calculate total irradiance; percent transmission was calculated as the total irradiance at 20 cm depth divided by the total irradiance at surface. Separately, to assess variation in spectral composition we first normalized the transmission spectra (integral equal to 1) to account for variations in brightness at different sites and dates.

This normalization allowed for our analysis to focus solely on variations in the light spectrum due to absorbance in the water column. We adapted a measure from Wilwert et al. (2021) and calculated the ratio of transmittance in the red (550-650 nm) vs. blue portions (350-450 nm) of the spectrum (Fig. S1). All calculations were completed in R (Team, 2021) utilizing packages Pavo (Maia et al., 2019) and Tidyverse (Wickham et al., 2019).

### Retinoid Extraction and Analysis

To determine the retinoid content of the red shiner eyes, we used High Performance Liquid Chromatography (HPLC). Retinoids were extracted from the whole right eye (left eye if right eye was not available) by grinding in 0.09% NaCl solution with 0.1 g of 2 mm zirconium beads (Next Advance, ZROB20) at 3000 hz for 90-120 seconds on a BeadBug Microtube Homogenizer (Model D1030).

This homogenization step was repeated two to three times for each sample. To derivatize the retinaldehydes we then added 400 µl of 2M hydroxylamine (Sigma, 255580) in distilled deionized water and 800 µl of methanol and incubated for at least 10 minutes at room temperature in the dark (Kane and Napoli, 2010). We then added 800 µl of acetone and 1.75-2 ml of hexane for retinaldehyde extraction. Samples were subsequently centrifuged for 3-5 minutes. The resulting solvent fraction was collected and dried under a stream of nitrogen. We then resuspended the extract in 120 µL of hexane, and injected 100 µL of sample into an Agilent 1100 series HPLC fitted with a Zorbax RX-SIL column (4.6x25 mm, 5 µm, Agilent). Elution of samples was achieved by a gradient mobile phase of 0.5% ethyl acetate in hexane for 5 minutes then increased to 10% ethyl acetate in hexane for 20 minutes. Isocratic conditions for 35 minutes followed. Throughout the run, the flow rate was 1.4 ml min_-1_ and the temperature of the column remained at 25°C. A photodiode array detector monitored the samples at 325, 350, and 380 nm. Authentic standards or published accounts were used to identify vitamin A_1_ and vitamin A_2_-derived retinoids (Kane and Napoli, 2010; Landers and Olson, 1988; Morshedian et al., 2017; Zonta and Stancher, 1984). Retinoid mass was calculated based on external standard curves for retinol and derivatized retinal. We summed the mass of all A_1_ or A_2_-derived retinoids including retinols, retinaldehydes, and retinyl esters and calculated the proportion of vitamin A_2_-derived retinoid as the A_2_ retinoid mass divided by total retinoid mass of the sample.

### Statistics analyses of light environment and chromophore composition

We carried out all statistical analyses in R (Team, 2021) using the base stats package except where noted. We compared turbidity among the site and seasons by fitting the linear model: `Log`_10_`(turbidity) ∼ site * season` and tested the effects of the independent variables with an ANOVA with type III sum of squares from using `car` package (Fox and Weisberg, 2018). For post hoc pairwise comparisons we estimate marginal means with the `emmeans` package (Lenth et al., 2019) and computed adjusted P-values by the Tukey method. We follow the same approach to compare our direct measures of spectral composition and a similar approach for light transmittance but fitted these proportional measures to a beta-regression model with `betareg` package (Cribari-Neto and Zeileis, 2010). To test for site and season effects on the relative abundance of A_2_ in the eyes of red shiners we fitted a beta-regression model: `A2 proportion ∼ site * season + eye diameter`. A number of individuals had A2 proportions = 0, which violates the assumptions of beta-regression. Therefore, we transformed all of the measures following the methods of (Cribari-Neto and Zeileis, 2010; Smithson and Verkuilen, 2006). We included eye diameter in the model because it varied nearly 2-fold among our samples and is an important determinant of the light gathering capacity of the eye (Cronin et al., 2014). We did not include sex as an independent variable because we were not able to determine the sex of all individuals we sampled. However, we analyzed a subset of individuals sampled in summer when nuptial coloration and gonad development allowed us to determine sex, and found that there were no significant differences in A_2_ abundance by sex (*F*_1,80_ = 0.75, *P* = 0.39) or the interaction of sex and sampling site (*F*_2,80_ = 0.12, *P* = 0.88). We calculated post hoc contrasts with `emmeans` as above and applied a Sidak adjustment (Lenth et al., 2019). Statistical analysis of gene expression is detailed below.

### RNA Extraction and Transcriptome Analysis

We extracted and purified total RNA from the left eyes of 16 red shiner samples. We selected four individuals each from low (Lightning creek) and high turbidity (Polecat creek) habitats in summer and late fall/winter. Each eye was transferred to a screw-cap tube with 1 ml of TRIzol reagent (ThermoFisher Scientific) and 0.1 g of 2 mm zirconium beads. We homogenized the whole eyes with a BeadBug Microtube Homogenizer (Model D1030) at 4 kHz for 180 seconds. We then extracted RNA following the manufacturer’s protocol with the addition of 1 µL of glycogen (R0551, Thermo Fisher Scientific) to facilitate RNA precipitation. To remove residual DNA, we treated the extracted total RNA with Turbo DNase (AM1907, ThermoFisher Scientific) following the manufacturer’s guidelines. We extracted the DNAse-treated RNA by adding 150 µL molecular grade water and 200 µL chloroform and mixing the samples by vortexing. We then centrifuged the samples, collected the aqueous fraction into a new tube, added 17.5 µL NaAC (3M pH 5.2), 1 µL glycogen, and 600 µL EtOH, and samples were incubated for 20 minutes at -20°C. Next, samples were centrifuged at 13,000 rcf for 10 minutes at 4°C, and the resulting supernatant was removed. Pellets were washed twice with 80% ethanol and air dried. Finally, the pellets were resuspended with 25 µL molecular grade water, and RNA quality and amount were measured utilizing the NanoDrop 8000 (Thermo Scientific).

The total RNA samples were sent to the Clinical Genomics Laboratory at the Oklahoma Medical Research Foundation (OMRF) for mRNA prep and sequencing. mRNA sequencing libraries were prepared by OMRF with the xGen RNA Lib Prep Kit (Integrated DNA Technologies) and NEB poly-A selection kit (New England Biolabs). The mRNA libraries were then sequenced as 150 bp paired-end reads on an Illumna NovaSeq 6000. We received demultiplexed reads in Fastq format and confirmed the quality of the raw reads with FastQC (v0.11.5) (Andrews, 2015). We then removed adaptor sequences and low-quality bases (Phred score <5) with Trim Galore! (v0.6.0) (Krueger, 2019).

### De Novo Transcriptome Assembly

A reference transcriptome was not available for *C. lutrensis*; therefore, we generated a *de novo* transcriptome assembly with Trinity (v2.8.4) (Grabherr et al., 2011) from the trimmed paired reads of two samples: a clear water summer sample (LCV_7_24Jul20) and a turbid water late fall/winter sample (PCJ_9_20Dec20). We supplemented the de novo assembly by referencing published red shiner opsin gene sequences by Chang and Yan (2019) and the *CYP27C1* sequence from this study. We used blastn (v2.13.0) (Camacho et al., 2009) to search the *de novo* assembly against the published opsin sequences and the *CYP27C1* sequence. We then merged the *de novo* and published sequences to create more complete opsin and CYP27C1 transcript sequences that included the 3’ and 5’ untranslated regions (Table S1).

### Analyses of selected transcripts

We pseudoaligned trimmed sequence reads in FastQ format to the supplemented *de novo* transcriptome assembly (described above), and generated read counts with Kallisto (v.0.46.2) using 50 bootstrap samples parameter `–b 50` (Bray et al., 2016; Melsted et al., 2019) (Table S2). We exported the counts for all genes as transcripts per million (TPM) using Sleuth (v.0.30.0) (Pimentel et al., 2017) and then extracted the count data for the opsins and *CYP27C1* for further analysis. We fitted separate linear models for each of the seven genes: `Log`_2_`(TPM) ∼ site * season` and tested the effects of the independent variables with an ANOVA with type III sum of squares, as described above. To account for multiple comparisons, we adjusted the P-values with the `p.adjust` function of base R and applied a Benjamini & Hochberg correction for seven comparisons. We also calculated the expression of each opsin as a proportion of total opsin expression of all six opsins in each sample and fitted a beta-regression model, tested with an ANOVA with type III sum of squares, and adjusted P-values as described above.

### Cloning and Functional Characterization of CYP27C1

To examine the catalytic function of CYP27C1 in the red shiner, we extracted RNA from the left eye of an adult female as described above. We generated cDNA by reverse transcription using an oligo (dT) 20 primer and superscript IV reverse transcriptase (18090010, ThermoFisher Scientific) following the manufacturer’s protocols. We amplified the full-length coding sequence of CYP27C1 by polymerase chain reaction with the forward primer: ataccgcaccggtagccaccATGGCTCTTCAAAGTACTATTCTACACATGG and reverse primer: ataccgcgcggccgcTTTTCGGTCTGTAAATCTAAGGTTGATGG. The lower-case sequence is additional sequence which contains the restriction sites for cloning. We digested the PCR products with AgeI and NotI (New England Biolabs) and cloned them into the first position of the bicistronic vector pCAG-[first position]-2A-GFP. We confirmed the CYP27C1 sequence with Sanger sequencing (Eurofins Genomics). We assayed enzymatic activity in HEK293 cells (ATCC, CRL-1573). Following the ATCC protocols, we cultured the cells to 80% confluency in a 6-well plate (8.96 cm_2_ wells) and then transfected with the CYP27C1 construct or a control construct (pCAG-dsRed-2A- GFP) using polyethylenimine (PEI, 23966-2, Polysciences, Warrington, PA). After 24 hours, we confirmed expression by visualizing GFP. We then dissolved 5 µg of retinol (Sigma, R7632) in 36 µl of ethanol and combined with 6 ml of complete media. We replaced the media in the wells with 1 mL each of the retinol-enriched media and incubated overnight (∼24 hrs). We then scraped the cells, discarded the media, and extracted carotenoids by adding 200 µl of distilled water, 200 µl of 100% ethanol, and disrupted the cells with 0.1g of 1 mm zirconium beads (Next Advance, ZROB10) at 4 kHz for 30 seconds on a Beadbug homogenizer. We then added 1 ml of hexane:tert-butyl methyl ether (1:1, vol:vol), homogenized again for 30 seconds, then centrifuged at 10,000 RPM for two minutes. We collected the upper solvent fraction from the sample and dried the extract under a stream of nitrogen. We then redissolved the cell extracts in 120 µl of hexane and analyzed retinoid content by HPLC, as described above.

## Results

### Light environment differs among study sites

We measured downward irradiance at 20 cm depth at each site and most sampling bouts and determined the transmittance relative to surface illumination and spectral composition. Light transmittance differed significantly among the sites by season (`site*season`: *F*_6,87_ = 13.008, *P* = 1.90 x 10^-10^, Fig. 1c). Post hoc analyses indicated that Big Creek had significantly greater transmittance than the other sites, in spring (Sidak adjusted comparisons *P* < 0.0022, Fig 1c). In summer, all three sites differed significantly (Sidak adjusted comparisons *P* < 0.0031) with the greatest transmittance at Lightning creek and lowest at Polecat creek (Fig. 1c). In the fall, Big Creek and Lightning Creek did not differ significantly, however, both had significantly greater transmittance than Polecat Creek (Sidak adjusted comparisons *P* < 0.0001, Fig 1c). In winter, only Big Creek and Polecat Creek differed significantly (Sidak adjusted comparisons *P* = 0.014, Fig 1c). Consistent with the historical turbidity measures (Fig. 1b), Lightning Creek tended to have the greatest transmittance (i.e. clearest waters), Big Creek was intermediate to the two other sites, and Polecat Creek had the lowest transmittance of all three sites. A notable exception to this trend was during spring, when we observed very low transmittance in Lightning Creek following heavy rainfall.

The spectral composition of the light environment differed significantly among sites by season (site*season: *F*_6,88_ = 39.592, *P* = 2.20 x 10^-16^, Fig. 1d). Post hoc comparisons indicate that the red to blue ratio differed significantly between Big Creek and Lightning Creek only during spring (Sidak adjusted comparison *P* < 0.0001).

However, during all seasons, the spectrum at Polecat Creek was significantly redder than the other sites (Sidak adjusted comparisons *P* < 0.0001, Fig. 1d). At Lightning Creek and Polecat Creek the spectrum was significantly redder in spring compared to other seasons (Sidak adjusted comparisons *P* < 0.0001, Fig. 1d). At Polecat Creek the spectrum was significantly blue shifted in the fall compared to all other seasons (Sidak adjusted comparisons *P* < 0.0001, Fig. 1d).

*Red Shiner Ocular Retinoid Composition:* Our initial HPLC profiling confirmed that the red shiner is a dual visual pigment species. We observed both A_1_ and A_2_ retinyl esters, aldehydes, and alcohols (Fig. S3) in the eyes of many of the individuals we sampled.

Retinal aldehydes were the most abundant of these forms in the eye (Fig. S3). Next, we investigated how retinoid composition varied among populations from different sites and seasons.

### A_2_-Derived Chromophore varies among sites and seasons

To understand if and how visual pigment chromophore composition varied, we sampled fish from different locations throughout the year and measured the retinoid content by HPLC. We found that the proportion of A_2_-derived chromophore differed significantly among sites by season (`site * season`: F_6,202_ = 3.0511, P = 0.0070, Fig. 2a). In all seasons, fish sampled from Polecat Creek, the most turbid site, had a significantly greater proportion of A_2_ chromophore than the other two sites (Sidak adjusted comparisons *P* < 0.0077, Fig. 2a). At each site, the proportion of A_2_ chromophore in the eyes of the fish differed significantly among the seasons with significantly lower proportions in summer compared to the winter samples (Sidak adjusted comparisons *P* < 0.0002, Fig. 2a). Eye size was also a significant predictor of A_2_ chromophore abundance (F_1,202_ = 5.7146, P = 0.018) and fish with larger eyes tended to accumulate a greater proportion of A_2_ chromophore (Fig. 2b)

**Figure 2.**
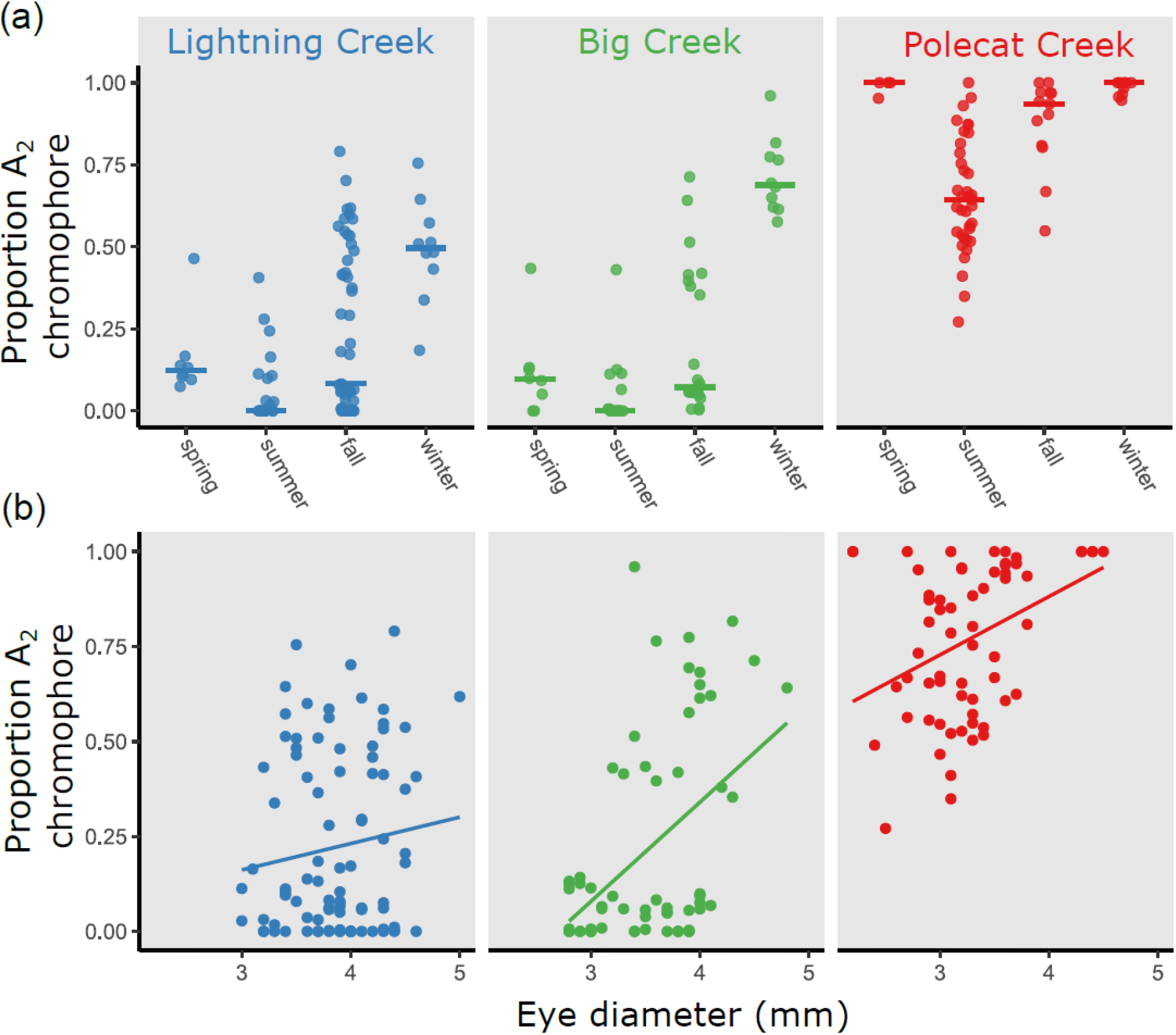
The chromophore composition of the red shiner eye differs among sites and with seasons and is correlated with eye size. (a) Proportion of A_2_ chromophore relative to total retinoid content of whole eyes. Each point represents an individual and the bar represents the mean proportion of A_2_ chromophore at each site in each season. (b) Proportion of A_2_ chromophore relative to eye diameter. Each point represents an individual and the line is a simple linear regression fit to the data from each site.

### CYP27C1 from Cyprinella lutrensis Catalyzes the Conversion of Retinol to 3,4- didehydroretinol

The enzyme CYP27C1 catalyzes the conversion of retinol to 3,4-didehydroretinol in zebrafish and evidence suggests that this mechanism is widely conserved among vertebrates (Enright et al., 2015; Morshedian et al., 2017). To confirm that the red shiner homolog of CYP27C1 possesses this same catalytic function, We cloned *C. lutrensis* CYP27C1, heterologously expressed the enzyme, and assayed its activity with a retinol substrate. The amino acid sequences of red shiner CYP27C1 is similar to the zebrafish homolog with 488/540 (90%) amino acid identities (Fig. S4). HEK293 cells expressing red shiner *CYP27C1* and supplemented with retinol produced a novel product that has a red-shifted UV-Vis absorbance spectrum and HPLC retention times consistent with 3,4- didehydroretinol (Fig. 3).

**Figure 3.**
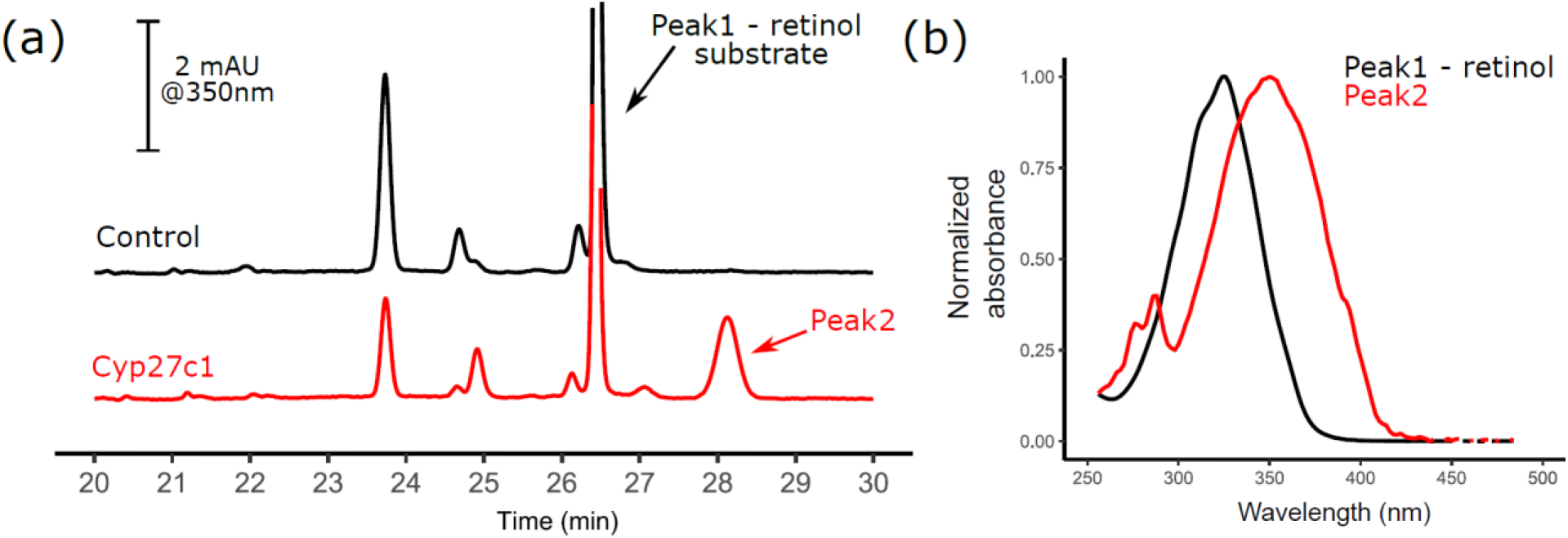
Red shiner CYP27C1 converts retinol to 3,4-didehydroretinol. (a) HPLC-DAD profiles of retinoids extracted from HEK293 cells transfected with a control construct (black) or red shiner CYP27C1 expression vector (red). (b) UV-Vis absorbance spectra of the retinol substrate (peak 1) and the novel product of CYP27C1 (peak 2). The retention time and UV-Vis spectrum of peak 2 is consistent with 3,4- didehydroretinol.

### Opsin and CYP27C1 gene expression patterns

Previous studies have demonstrated adaptive plasticity in opsin expression is linked to light environment and that *CYP27C1* is the enzyme that catalyzes the conversion of A_1_ to A_2_ derived chromophore (Carleton et al., 2020; Chang and Yan, 2019; Corbo, 2021). Therefore, we wanted to know if and how the expression of these genes varied in the eyes of red shiners sampled from our study sites. To do this, we sequenced the ocular transcriptomes of a subset of individuals from the most (Polecat Creek) and least (Lightning Creek) turbid sites in summer and fall/winter seasons. We sequenced the transcriptomes to an average depth of 24,114,599 reads per sample with an average of 52.23 percent alignment with our *de novo* transcriptome assembly (Table S2).

In contrast to the dramatic differences in A_2_ chromophore abundance among sites and across seasons (Fig. 2a), we did not observe significant differences in *CYP27C1* expression among our subset of samples (`site`: F_1,12_ = 4.00, *P_adj_* = 0.16, `season`: F_1,12_ = 0.28, *P_adj_* = 1.00, Fig. 4a). *CYP27C1* expression ranged from 0 to 9.2 TPM among the samples, a level of expression that is generally considered “low” (Papatheodorou et al., 2018).

**Figure 4.**
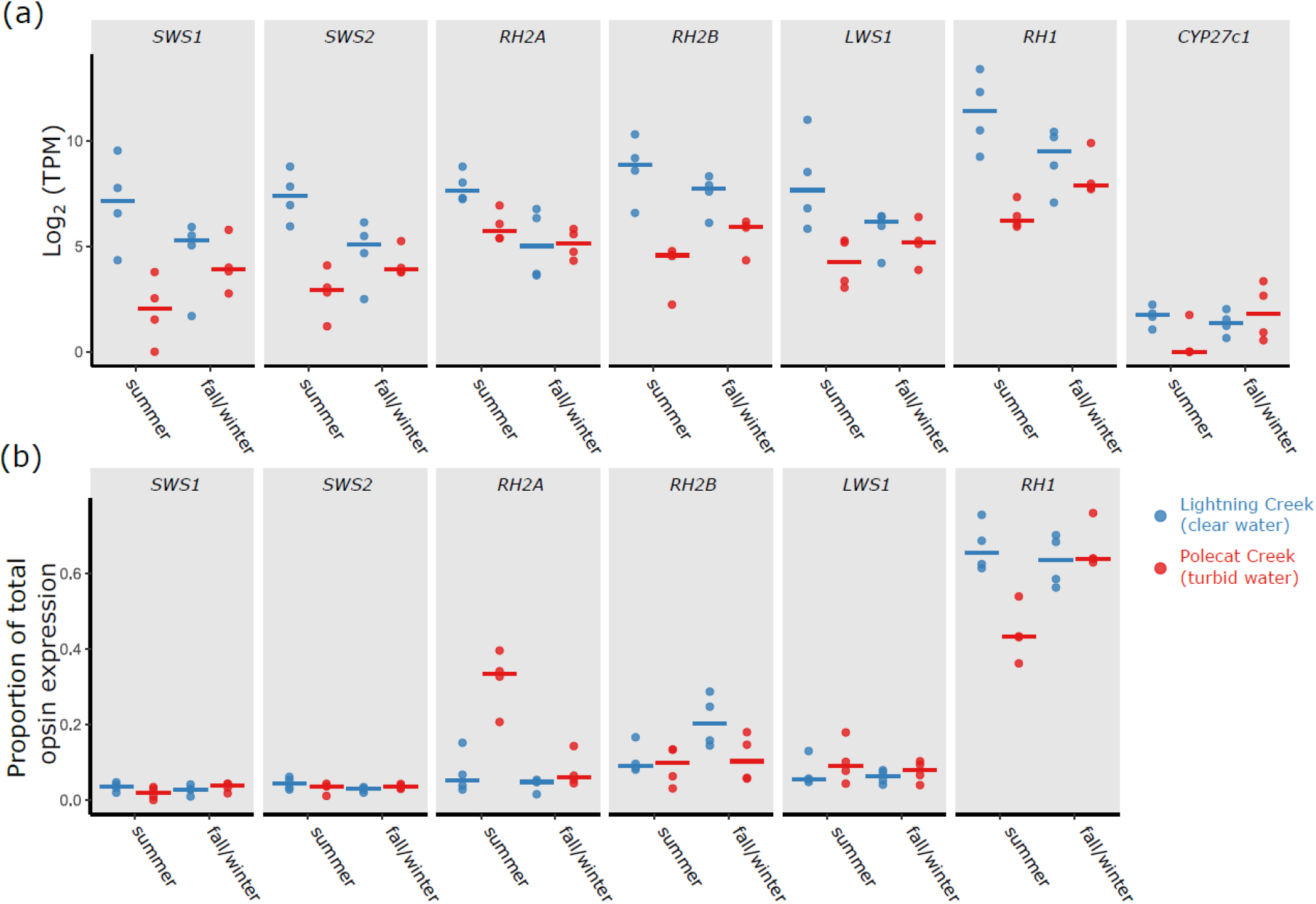
*Opsin* and *CYP27C1* expression. (a) Normalized gene expression measured as transcripts per million (TPM) for individual samples at a clear water site (Lightning Creek) or turbid water site (Polecat Creek). Each point represents an individual and bars indicate mean expression at each site in each season. (b) The expression of each opsin gene relative to total opsin expression with an individual eye. Each point represents an individual and bars indicate mean relative expression at each site in each season.

We compared the expression of the opsins two different ways. First, we directly compared the normalized transcript counts of each opsin (transcripts per million – TPM) and then, to capture possible opsin-based shifts in spectral sensitivity, we calculated the proportional expression of each opsin relative to the total opsin expression in the sample. *Rhodopsin* 1 (*RH1*) was the most highly expressed opsin in all samples (Fig 4) and the normalized and relative expression of *RH1* differed significantly between sites by season (`TPM ∼ site * season`: F_1,12_ = 9.45, *P_adj_* = 0.023, `proportion ∼ site * season`: F_1,11_ = 18.37, *P_adj_* = 0.003). The relative expression of *RH1* in the Lightning Creek (low turbidity site) sample was significantly greater than the Polecat Creek (high turbidity site) samples in the summer season (Sidak adjusted comparisons *P* < 0.0001, Fig. 4b), and relative *RHO* expression in the Polecat Creek samples significantly increased from summer to the late fall/winter samples (Sidak adjusted comparisons *P* < 0.0001, Fig. 4b).

*Long wavelength-sensitive opsin 1 (LWS1)* normalized expression was significantly greater in the Lightning Creek samples (site: F_1,12_ = 13.51, *P_adj_* = 0.011), but there were no significant differences between sites or seasons in *LWS1* relative expression (Fig. 4b).

Like many other teleost fish, the medium wavelength sensitive *rhodopsin 2 (RH2)* has been duplicated, and two forms, RH2a and RH2b, are present in the red shiner genome (Chang and Yan, 2019; Musilova and Cortesi, 2023). The normalized expression of the *RH2a* differed significantly between seasons with higher levels observed in the summer samples (`season`: F_1,12_ = 13.54, *P_adj_* = 0.011, Fig. 4a). The relative expression of *RH2a* differed significantly among sites (`site`: F_1,11_ = 36.53, *P_adj_* = 0.00029) with levels significantly higher in the Polecat Creek (high turbidity) samples in the summer (Sidak adjusted comparisons *P* < 0.0001) and declining significantly from summer to late fall/winter (Sidak adjusted comparisons *P* < 0.0001). The normalized expression of *RH2b* was significantly greater in the Lightning Creek samples (`site`: F_1,12_ = 31.00, *P_adj_* = 0.0004), but there were no significant differences in relative expression of *RH2b* between sites or seasons (Fig. 4b).

The normalized expression of the *short wavelength-sensitive opsins 1* and *2* (*SWS1* and *SWS2*) differed significantly between the sites by season (*SWS1* `site * season`: F_1,12_ = 6.81, *P_adj_* = 0.053, *SWS2* `site * season`: F_1,12_ = 11.34, *P_adj_* = 0.013, Fig. 4a). The normalized expression of *SWS1* and *SWS2* was significantly greater in Lightning Creek (low turbidity site) samples in the summer (Sidak adjusted comparisons *P* < 0.0064, Fig. 4a) and *SWS2* levels in Lightning Creek samples decline significantly from summer to late fall/winter (Sidak adjusted comparison *P* < 0.035, Fig. 4a). There were no significant differences in *SWS1* or *SWS2* relative expression between sites or seasons (Fig. 4b).

## Discussion

Our results show that the red shiner is a dual chromophore species with a plastic visual system where the relative abundance of the A_1_ and A_2_ visual pigment chromophores differed significantly among habitats and seasons. We demonstrate that red shiner *CYP27C1* catalyzes the conversion of vitamin A_1_-derived chromophore to vitamin A_2_-derived chromophore, but the expression patterns of *CYP27C1* in the eye are not consistent with the patterns of A_2_ chromophore abundance. We observed significant variation in visual pigment opsin expression among habitats and seasons, but this variation was not consistent with patterns of environmental variation reported in other teleost fish.

### Chromophore composition matches median light environment and shifts with the seasons

We predicted that red shiners inhabiting turbid waters would have a higher proportion of A_2_ chromophore in their eyes than those inhabiting less turbid water. Consistent with this prediction, we found the proportion of A_2_ chromophore to be significantly greater in fish from the historically most turbid site, Polecat Creek, compared to the sites with clearer waters, Lightning Creek and Big Creek. This pattern of turbidity-dependent chromophore composition is consistent with studies of the golden shiner (*Notemigonus crysoleucas*) and a variety of cichlid species that show greater proportions of A_2_ chromophore in the eyes or photoreceptors of individuals sampled from turbid habitats (Bridges, 1964; Carleton and Yourick, 2020; Escobar-Camacho et al., 2019; Härer et al., 2018; Terai et al., 2017). These habitat-specific patterns of chromophore composition are considered adaptive, and it is hypothesized that the a shift from an A_1_ to A_2_ chromophore dominated retina will red-shift visual sensitivity to match the red-shifted light environment of turbid waters (Bridges, 1964; Bridges, 1972; Corbo, 2021; Enright et al., 2015; Escobar-Camacho et al., 2019). Consistent with this hypothesis, our measures confirmed that the available light spectrum in the most turbid site, Polecat Creek, was significantly red shifted compared to the clearer water sites.

The light environments at our study sites were dynamic, and we observed considerable variation within seasons and significant changes across the seasons. Laboratory investigations of the A_1_ to A_2_ chromophore switch in zebra fish (*Danio rerio*) indicate that these changes occur over the course of weeks (Enright et al., 2015).

Therefore, short-term changes in light environment, like the influx of turbid water following a single thunderstorm, are unlikely to be closely tracked by changes in chromophore composition. To test this prediction, we refit our model predicting A_2_ chromophore with our direct measures instead of site as a factor, compared the fits of these models by AICc, and found that these models were poorer fits (delta AICc > 102) than the “site” model (Table S3). This result suggests that the mechanisms of chromophore plasticity may be integrating over a longer-time scale and tuning the visual system to the mean or median conditions in a habitat.

We observed seasonal changes in chromophore composition, but our results suggest that these are not strongly linked to the light environment. Historical turbidity measures indicate that turbidity tends to be lower in the winter months and peaks during the rainy late spring and early summer season. Our direct measures of light environment are partially consistent with this trend, with some of the sights showing increased transmittance and a relative blue shift in the spectrum. Therefore, if red shiners are matching their visual sensitivities to these seasonal changes in the light environment, we expect a shift from the A_2_ chromophore to the relatively blue-shifted A_1_ chromophore during the winter months. However, this prediction was not supported, as the proportion of the A_1_-derived chromophore decreased in the winter months in the eyes of red shiners at all sites. This suggests that chromophore composition is responding to something other than the spectral composition of light.

A shift toward increased A_2_ chromophore levels in the winter months has been observed in a diversity of fish species and also an amphibian (Bridges, 1964; Bridges, 1965; Bridges, 1972; Makino et al., 1983; Muntz and Mouat, 1984; Ueno et al., 2005). Detailed experimental investigations in the 1970’s and 80’s identified temperature and light as important drivers of chromophore composition. The effects of experimental light manipulations are complex. Total darkness, occlusion of the eye, or reduced daylight hours consistently favor a shift to and A_2_ chromophore dominance (Allen, 1971; Allen and McFarland, 1973; Bridges and Yoshikami, 1970; Tsin and Beatty, 1977), but there is also evidence that high light intensity leads to increases in A_2_ chromophore levels (Allen, 1971; Cristy, 1976). In contrast, across all studies, low temperatures consistently resulted in a shift from A_1_ to A_2_ as the dominant chromophore in the eye (Allen and McFarland, 1973; Cristy, 1976; McFarland and Allen, 1977; Tsin and Beatty, 1977).

Seasonal and temperature driven shifts in chromophore composition may reflect an adaptive balance of spectral tuning and photoreceptor noise. A_2_ containing visual pigments are more susceptible, than A_1_ visual pigments, to spontaneous isomerization in the absence of light (Ala-Laurila et al., 2007). Therefore, the shift from an A_1_ to A_2_ chromophore increases dark noise within the cell and reduces the sensitivity of the photoreceptor (Ala-Laurila et al., 2007; Barlow, 1956). A_2_ containing visual pigment instability is positively temperature-dependent however, and this dark noise is predicted to be less at lower temperatures (Aho et al., 1988). Therefore, the cost (loss of visual sensitivity) may be reduced in cold conditions as red shiners are able to utilize A_2_ chromophore and expand the light spectrum that is visible without the cost of spontaneous isomerization. Despite these seasonal shifts, chromophore differences among the study sites remain, suggesting the regulation of chromophore composition is finely tuned and likely integrates several environmental cues. Thus, the red shiner offers an excellent opportunity to deconstruct these mechanisms of regulation and a readily accessible natural system to investigate how local environments drive the evolution of this plasticity.

### CYP27C1 expression patterns are inconsistent with chromophore composition

The enzyme CYP27C1 has been identified as necessary and sufficient to catalyze the conversion of vitamin A_1_ to vitamin A_2_ in zebrafish (Enright et al., 2015). Subsequent studies suggest that this mechanism of chromophore metabolism is widely conserved among vertebrates (Morshedian et al., 2017). Consistent with this conserved function, we found that red shiner CYP27C1 was sufficient to convert retinol (A_1_) to 3,4- didehydroretinol (A_2_) in our cell culture assay system. Therefore, we hypothesized that changes in *CYP27C1* expression were mediating the environmental and seasonal variation and predicted that *CYP27C1* expression levels in the eyes of red shiners would track A_2_ chromophore abundance. However, this prediction was not supported by our transcriptome profiling analyses of whole eyes. We found that *CYP27C1* expression was low in all samples and there were no significant differences in expression among sites or seasons. However, this result does not rule out a role for CYP27C1 in chromophore conversion. We extracted RNA from whole eyes, therefore our ability to detect changes in *CYP27C1* may have been confounded by the complex collection of tissues and cell types in the samples. Enright et al. (2015) found that CYP27C1 expression concentrated in the retinal pigment epithelium of the eye, and a targeted analysis of these cells may reveal a different pattern. Vitamin A_2_ has been found to be the dominant retinoid form in the plasma and liver tissues of several fish species, suggesting that the conversion of vitamin A_1_ to vitamin A_2_ might be occurring outside of the eye (Balasundaram et al., 1956; Defo et al., 2012; Goswami and Barua, 1981). An investigation of the distribution of A_1_ and A_2_ dynamics in other tissues of the red shiner is warranted.

The ocular expression of *CYP27C1* has been used as a proxy measure for chromophore composition in several recent studies (Karagic et al., 2022; Wilwert et al., 2022; Wilwert et al., 2023). However, in our study, patterns of chromophore abundance and ocular expression of *CYP27C1* were not concordant. A similar discordance was observed by Escobar-Camacho et al. (2019) where they found that ocular *CYP27C1* expression was significantly lower among turbid habitat fish with higher A_2_ levels. These results indicate that the assumed correlation between ocular *CYP27C1* levels and chromophore composition should be validated within each study system.

### Opsin expression varied among habitats and seasons

Chang and Yan (2019), in a laboratory study, demonstrated that opsin expression is plastic and that red shiners increase *LWS* opsin, and decrease SWS opsin expression in turbid, red-shifted light environments (Chang and Yan 2019). Therefore, we predicted that the red shiners we sampled from turbid environments would show a similar pattern of elevated *LWS* and reduced *SWS* opsin expression compared to those sampled from clearer water conditions. Contrary to this prediction, we found no significant differences in the relative expression of *LWS1, RH2b, SWS1*, or *SWS2* between the site or seasons. There are several reasons why our results might differ from Chang and Yan (2019). While the fish in the previous study were purchased from aquarium suppliers and reared under captive conditions, we studied wild-caught fish. The artificial light environments of the captive study may differ considerably from the habitats we sampled. The lowest turbidity environment in the lab study was 0 NTU, and the significant variance in *LWS* expression was found only between 0 NTU and each higher turbidity environment (50, 100 and 200 NTU) (Chang and Yan, 2019). The median turbidity at our clear water site was 3.0 NTU, and the turbid site 20.4 NTU. Perhaps the most turbid lightning environment (Polecat Creek) in our study was not challenging enough to elicit significant variation in opsin expression.

The only cone opsin that differed in its relative expression was the medium wavelength sensitive *RH2a* with relatively high expression among fish sampled in the most turbid conditions (summer, Polecat creek). Diverging response in the expression of the *RH2a and RH2b* opsin paralogs have also been observed among guppies (*Poecilia reticulata*), raising the interesting possibility that these opsins may be sub- functionalized in a yet unknown way (Ehlman et al., 2015; Musilova and Cortesi, 2023).

The relative expression of rod opsin *RH1* was significantly higher among fish from the clear water sites, and its expression among the turbid-site fish increased from summer to the fall/winter season. This suggested that the relative expression of *RH1* may be influenced by water clarity and positively correlated with available light levels. Among Lake Tanganyika cichlid species, *RH1* expression is positively correlated with eye size, an important determinant of the amount of light reaching the photoreceptors (Ricci et al., 2023). Rod photoreceptors mediate scotopic vision, therefore the shifts we see in the relative expression of *RH1* might reflect or impact diel activity patterns of red shiners in different habitats. In other words, are fish in clear water sites more active in low light conditions?

When we examined opsin transcript levels (TPM values), rather than expression relative to the other opsins, we found that the expression of most of the opsins was significantly higher in samples from the clear water site, especially in the summer season. This finding is contrary to previous studies findings where increased expression of cone opsins was observed in turbid conditions compared to clear water conditions (Chang and Yan, 2019; Fuller and Claricoates, 2011). The transcript level variations we observed are consistent with light levels and day-length driving opsin expression in general. For example, daily rhythms of opsin expression have been identified in the Senegalese sole and a species of African cichlid (Frau et al., 2020; Halstenberg et al., 2005). Sole kept under alternating light and dark conditions, with either white or blue light spectra, had peak opsin expression at the end of the light period or at the second half of the day (Frau et al., 2020). Similarly, Halstenberg et al. (2005) found all cone opsins analyzed peaked in expression during the late afternoon in the African cichlid, *Haplochromois burtoni*. Therefore, long days in the summer months and clear water conditions may promote opsin expression in general.

## Conclusions

The red shiner’s abundance, wide distribution in North America, and dual pigment visual system make it an excellent model organism for visual system plasticity research. We have demonstrated significant variation in visual pigment chromophore composition relative to light environment, and season suggesting this is a plastic trait. These variations in chromophore composition are likely to have a substantial impact on the sensitivities and function of the red shiner visual system and highlight the importance of considering the A_1_ to A_2_ chromophore switch as an integral part of visual system plasticity. The significant relationship between chromophore composition and season suggests there may be a trade-off between spectral tuning and receptor noise thresholds, presenting an exciting opportunity to understand how multiple environmental cues are integrated to shape plastic responses. Opsin expression has historically been the focus of visual system plasticity research. However, our results indicate that habitat and season related variation in chromophore composition is greater than variation in opsin expression. Therefore, in red shiners, and possibly other fish species, chromophore composition may be a more important mechanism of visual system plasticity.

## Supporting information

Supplemental Figures and Tables for Foster et al.

## Acknowledgments

We thank Matthew Dugas for his advice on red shiner sampling, study site selection, and study design. We thank Ronald Bonnet and Alexandra Kingston for their contributions to study design, analysis, and interpretation. We thank Dustin Smith and Desirae Gonzales for their assistance in the field. We are grateful to Lance Phillips at the Oklahoma Water Resources Board and Karla Spinner at The Oklahoma Conservation Commission for providing access to historical turbidity records. The computing for this project was performed at the OU Supercomputing Center for Education & Research (OSCER) at the University of Oklahoma (OU).

## Author contributions

Conceptualization: T.F., M.T.; Methodology: T.F., B.F., M.T.; Formal analysis: T.F., A.W., B.F., M.T.; Investigation: T.F., A.W., B.F., M.T.; Resources: M.T.; Writing - original draft: T.F., M.T.; Writing - review & editing: T.F., A.W., B.F., M.T.; Visualization: T.F., M.T.; Supervision: T.F., M.T.; Funding acquisition: M.T.

## Data availability

RNA sequencing data have been deposited with the NCBI short-read archive (BioProject ID: PRJNA1076457). The transcriptome assembly and other raw data files and analysis scripts are available in the Dryad Digital Repository - https://doi.org/10.5061/dryad.vhhmgqp2c - temporary reviewer link - https://datadryad.org/stash/share/J8mL6UfTw-gS0AUslijiN9UMw7nSYbMYkLKs0U7MbGQ.

