## Supplemental Figures and Tables for Foster et al. for "Light environment and seasonal variation in the visual system of the red shiner (*Cyprinella lutrensis*)"

Figure S1. (a) a map of the study sites and the location of the University of Tulsa main campus. River and lake data courtesy of the Oklahoma Water Resources Board. (b) Representative downwelling irradiance spectra at 20 cm depth from each site. The blue and red portions of the spectrum used in the calculation of the red to blue ratio are highlighted.

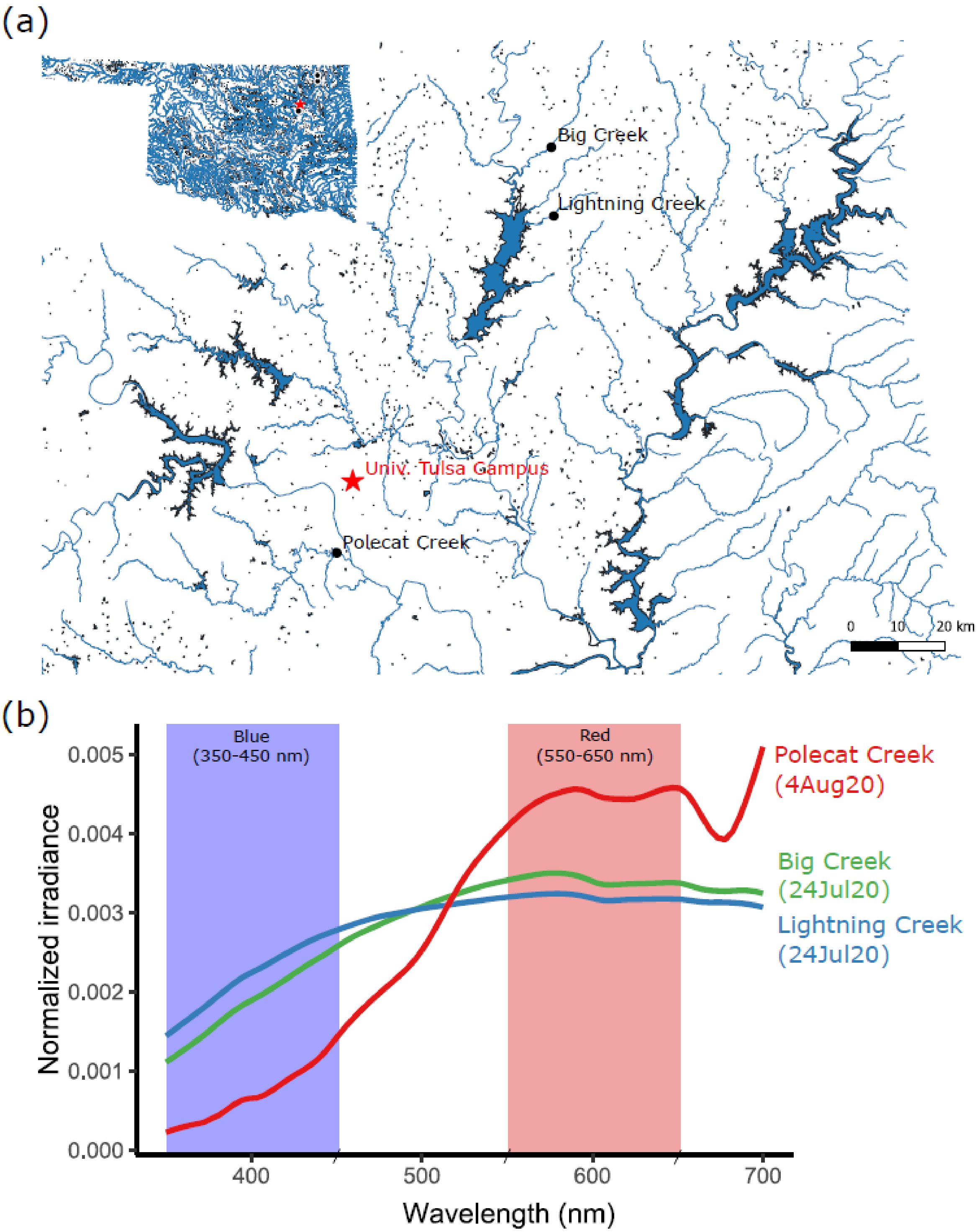

Table S1. Transcript sequences of the *C. lutrensis* visual opsins and *CYP27C1* assembled in this study.

```
>RH2A_DN1250_c0_g2_i7

GAGGAGGAAAGAGCAGATCCAACACTGCAGATCGCATCCTCTTCCAGGTCTGGATCACTAGTTGGCAAAGATGAACG
GCACTGAGGGAAACAACTTCTACATCCCCATGTCCAACAGGACAGGGCTAGCGAGGAGTCCTTTCGAATATCCACAG
TATTATCTTGCTGAGCCGTGGCAGTTTAAAGTGCTTGCTGTTTACATGTTCTTCCTCATCTGCTTGGGTTTTCCCAT
CAATGTCCTTACCTTGCTGGTTACAGCTCAACACAAAAAGCTCAGGCAACCTCTCAACTTCATTTTGGTCAACCTGG
CTGTGGCTGGCACCATCATGGTTTGTTTTGGATTCACGGTCACTTTCTACACAGCGATTCATGGCTACTTTGCTCTG
GGTCCAACTGCCTGCGCCATTGAGGGCTTCATGGCCACACTTGGAGGTGAAGTYGCCCTGTGGTCACTTGTGGTACT
GGCCATCGAGAGATACATTGTGGTTTGCAAGCCAATGGGTAGTTTCAAATTCTCAAGCACCCAYGCTTTGTCAGGAA
TTGGATTTACATGGGTAATGGCAATGGCATGTGCAGCTCCCCCTCTGGTTGGCTGGTCCAGATATATTCCTGAGGGA
ATGCARTGTTCATGTGGACCAGACTACTACACTCTGAGTCCTGGATACAACAATGAATCATATGTTCTCTATATGTT
CTCCGTCCATTTTATACTTCCGGTTACCGTAATCTTCTTCACCTATGGGCGGCTTGTTTGCACCGTCAAGGCGGCTG
CAGCTCAACAGCAGGACTCAGCATCCACCCAGAAGGCTGAGAGGGAAGTGACAAAAATGGTCATCCTGATGGTTTTC
GGTTTCTTGTTGGCTTGGACCCCTTATGCCACTGTTGCTGCTTGGATCTTCTTTAATAAGGGAGCTGCTTTCAGTGC
CCAGTTCATGGCTGTTCCTGCCTTTTTCTCAAAGAGCTCAGCCATATATAACCCTGTCATCTATGTGCTTCTAAACA
AACAGTTCAGGAACTGCATGCTGACCACWCTTTTCTGTGGAAAGAATCCTCTTGGTGATGAGGAGTCCTCAACTGTG
TCCACCAGCAAGACGGAGGTGTCCTCTGTATCTCCAGCATAGACTTTTGAACCTCTCACAGATTAAGTTCTTTTTGT
CATTGACCAAATTGGGGATCTGAAAAAAAGACCAAAAAAAAAGACAGTTGAATTAGTTTCTGCTTGCGTTCACTATC
CTATTATATGTATTTAGAGCTCATGGTTTATTGGGTGATCAGTGAACATTGAAATTGTGGTTTTATTTTTATGTGCC
GGGAGGCTAGCATTTATATCTTCTGCCGAAAATGTACATATATTTTGTAAATAATGAAAAAGAAATACATAGTAATA
AATACTGGCATTCAAAAAAAAAAAAAAAAAAAAA

>RH2B_DN1250_c0_g2_i6

GAGGAGGAAAGAGCAGATCCAACACTGCAGATCGCATCCTCTTCCAGGTCTGGATCACTAGTTGGCAAAGATGAACG
GCACTGAGGGAAACAACTTCTACATCCCCATGTCCAACAGGACAGGGCTAGCGAGGAGTCCTTTCGAATATCCACAG
TATTATCTTGCTGAGCCGTGGCAATTTAAACTGCTTGCTGTTTACATGTTCTTCCTCATCTGCTTGGGTTTTCCCAT
CAATGTCCTTACCTTGCTGGTTACAGCTCAACACAAAAAGCTCAGGCAACCTCTCAACTTCATTTTGGTTAACCTGG
CTGTGGCTGGCACCATCATGGTTTGTTTTGGATTCACGGTCACTTTCTACACAGCAATTCATGGCTACTTTGCTCTG
GGTCCAACTGCCTGCGCCATTGAGGGCTTCATGGCCACACTTGGAGGACAAGTTGCCCTTTGGTCACTCGTGGTACT
GGCCATCGAGAGATACATTGTGGTTTGCAAGCCTATGGGTAGTTTCAAATTCTCAACCACCCACGCTTTGTCAGGAA
TCGCATTTACATGGGTAATGGCCATGACATGTGCGGCTCCCCCTCTGGTTGGCTGGTCCAGATATATTCCTGAGGGA
ATGCAGTGTTCATGTGGACCAGACTACTACACTCTGAGTCCTGAATACAACAATGAATCATAYGTTTTCTACATGTT
CACCTGCCATTTTATAACCCCGGTTACCATAATCTTCTTCACCTATGGGCGGCTTGTTTGCACCGTCAAGGCGGCTG
CAGCTCAACAGCAGGACTCAGCATCCACCCAGAAGGCTGAGAGGGAAGTGACAAAAATGGTCGTCCTGATGGTTTTG
GGTTTCCTGGTGGCTTGGACCCCTTATGCCTCTGTTGCTGCTTGGATCTTCTTTAATAGGGGAGCTGCTTTCAGTGC
CCAGTTCATGGCTGTTCCTGCCTTTTTCTCAAAGAGCTCAGCCATATTTAATCCTATAATTTATGTGCTACTAAACA
AACAGTTCAGGAACTGCATGCTGACCACTCTTTTCTGTGGAAAGAATCCTCTTGGCGATGACGAGTCCTCAACTGTG
TCCACCAGCAAGACGGAGGTGTCCTCTGTATCTCCAGCATAGACTTAAAAAAAAAAACAAGGCGAATTAGGTGATGC
GTCTCATAAGGTTGTGTTGACAATGACTATTAAACTGGTGGACATACATTTGTTAAATAAGTAAAAGGCTTAAAGGG
ATAGTTCAGAAAATGAAAATTATCCCATAATTTACTCTCCCTCAGAAGGCCTTTACCCCCTGGAGCCATGGATAATC
TTATAATCTTAAATATCTGGCAATACTCGCTTCCATTATATAGCTTGGAAAAGGCAGGATATTTTAGAAACCTATAG
GATGGCTTGAGTGTGAGTAAATTGTGGGATAATTTTCCTTTTTGGGTGAATAACCTTTTAATGGGTTATCGGTGGTT
ATTTTTTATTAACTGGAAGGCTAACATTTATATCTTTGTCCTGAAATGTGTACATATTTTGTAAATAATAAAAAAAG
GATAGAAATAAATTCTGGCATTAAAAAAAAAAAAAAAAAAAAA

>SWS1_TRINITY_DN999_c0_g1_i2
```

GGGAGAGTCAGGCTGCTCGTTGACTTCACCAGGACACCGTGCTAGAACAAGGCCCCATACAGCGCAACCATGGACGC
ATGGAGCGTTCAGTTCGGGAACCTTTCCAAAGTCAGCCCCTTCGAGGGCCCACAGTACCACCTGGCCCCCAAGTGGG
CTTTCTACCTGCAGGCAGCTTTCATGGGCTTCGTATTTTTCGTGGGCACCCCTCTGAACGCCGTCGTCCTCTTCGTT
ACAATGAAGTACAAGAAGCTTCGACAGCCTCTCAACTACATCCTGGTGAACATCTCCCTAGCAGGCTTCATTTTCGA
CACGTTCTCTGTTAGCCAAGTATTCGTTAGCGCTCTTAGAGGTTATTACTTCCTCGGTCACACGTTATGCGCGTTGG
AAGCAGCAATGGGRGCTATTGCGGGTCTTGTGACGGGATGGTCCCTTGCCGTTCTGGCTTTCGAGAGATACGTGGTC
ATCTGCAAACCCTTCGGAGGCTTCAAGTTCGGACAAAGCCAGGCCTTGGGTGCCGTGGCGTTCACCTGGGTCATTGG
CACCGGCTGTGCCACTCCTCCATTCTGGGGATGGAGCAGATACATTCCCGAGGGTCTCGGCACCGCCTGCGGACCTG
ACTGGTACACAAAAAGCGAGGAGTATAATTCRGAGAGCTACACTTACTTCCTCATCGTCACCTGCTTCATCATGCCA
ATGACCATCATCATCTTCTCTTACTCACAGCTCTTGGGAGCCCTGCGTGCTGTTGCAGCCCAGCAAGCTGAGTCAGC
CTCTACCCAGAAGGCCGAGAAGGAAGTGTCCAGGATGGTTATTGTAATGGTCGGCTCTTTTGTGCTTTGCTATGCCC
CCTATGCCGCAACTGCTATGTGGTTTGCCAACGCTGATGAATCAAACAAAGATTACCGTCTGGTAGCCATCCCAGCT
TTCTTCTCAAAGAGCTCCTGCGTRTACAATCCCCTAATCTACGTCTTCATGAACAAACAGTTCAATGCCTGCATCAT
GGAGACAGTATTTGGCAAGAAGATTGATGAGGCTTCAGAGGTTTCCAGCAAGACCGAAACCTCTTCTGTGGCTGCAT
AAATTATTCCAGACCACGTTTAAAATTTCCCAGGACCTGTTTTCCTGTTATTCTCCCCCTGATTGTTCAAATCCCGT
ACCTTTACACATTGTCAAGGGCCAAGGGAGCCCCTTTGTCCCGAAGACACAGCCCACTGGATGGCCAAGCCGGGGGC
GATATCCAACTATCCTTTCTGAACTGGTACCTCTTCAACTACATCTTTTCTTTATTTTTTTGTAATGAAAAACATTG
CTATTTCATGTTAAAACAATGCATTATGCATTGATGATGAAGTAGAATATTTAATTAACTTGCAAAAGAAAATATAA
AAAAAAAAAATGATAATAATTGATAATGGAGAATCTCTTGTGTGAATGGACTCAGGCAAACTGAAAATGGAAATTCC
TGGCTTTTGTTTTTAAAATGCTCTTTGACTTACCCATTAGGCACGATGCACTATAATTATGTTTTTTATTCAATCAC
AATGAGAGCTGAAGGAAACTAAGCATTGAGAATATTCCAACTCTCCATGTCAATCAAAAGTACTTATTTCGACAAAA
GAGCAGAATTTAACTAAGCGGTTTACTCTCCTCTCCTCTAAAGAACCTACTCAGGCCCAGCTTTCTGGCTGGCTTAG
CCAAGATAAGTAAGATCACAAGGAATTATCCGAGTCCATCCGCTTGTTTGCAGCTGAGGAACCTTTCATTGGAAGTT
TGCTGGTAACAAATAATGAGCGACAGCTCTTGTCCAATCACTGGCTTCATGACGTTGGATCTGTAATATCTGTATCT
ATATTACCTCTGTACATGTTCTGTTGTAATTAGACATGCAGTGTGCATGAATTTGAACATGTTCACTGTTTTTTAGA
ATTAATAAAGATTTATATTGTTTGATGCAAAAAAAAAAAAAAAAAAAAAAAAAAAAATTAAATTATATATAGGAAAAT
CACACATCTTAACTACAATCACAGTTACATTTATATCAT

>LWS1_TRINITY_DN720_c0_g1_i8

AGGGAAGGTCTGTGCGAGTCGGCCGGAGACAGAAGGTAGTAAAAACACAAAGGCCCAAAGCTAAGTGACTACAGGTT
TTGGGCTATACAACATCCTTCAAAAATGGCAGAGCATTTGGGAGAAGTGATGTTTGCAGCCAGGCGAAGAGGAGAGG
AAACAACAAGAGAAGCAATGTTCACATATACCAACAGCAATAACACCAAAGATCCCTTTGAGGGACCCAACTACCAC
ATTGCCCCTCGATGGGTGTACAACGTCTCAACATGCTGGATGATATTTGTAGTAATTGCTTCGGTCTTCACCAATGG
CCTGGTGCTGGTTGCCACGGCCAAATTCAAGAAGCTCCGTCACCCCCTCAACTGGATTTTGGTCAACCTCGCTATAG
CTGATCTAGGAGAGACAGTGTTGGCCAGCACCATCAGTGTGATCAACCAGATTTTCGGCTACTTTATCCTCGGACAT
CCCATGTGTGTCTTTGAGGGCTACACTGTATCTGTATGTGGTATCTCTGCTCTGTGGTCTTTGACTATCATCTCTTG
GGAGAGATGGGTGGTCGTCTGCAAACCATTTGGAAATGTCAAGTTTGATGGTAAATGGGCAGCTGGTGGCATCATCT
TCTGCTGGGTTTGGTCTGCTGTCTGGTGTGCACCTCCCATCTTTGGCTGGAGCAGATATTGGCCTCAYGGTCTGAAG
ACCTCTTGCGGCCCTGATGTCTTCAGTGGAAGCGAGGACCCCGGTGTCCAGTCCTACATGATTGTCCTAATGATCAC
CTGCTGTATCATACCTCTGGCCATCATCATTCTCTGCTACATTGCWGTGTACTTGGCCATTCATGCTGTCGCCCAGC
AACAGAAGGATTCTGAGTCCACACAGAAAGCTGAGAAGGAAGTGTCCAGAATGGTGGTTGTCATGATCGTGGCCTAC
TGTGTTTGCTGGGGTCCTTATACGGTCTTTGCCTGCTTTGCAGCTGCAAACCCAGGCTATGCCTTCCATCCACTGGC
AGCAGCCATGCCTGCCTACTTTGCCAAGAGCGCCACCATCTACAACCCCGTCATTTACGTCTTCATGAACCGACAAT
TCCGTGTATGCATCATGCAGCTCTTTGGAAAGAAGGTGGATGATGGGTCTGAGGTGTCTACATCCAAGACAGAAGTG
TCCTCTGTGGCTCCTGCATAAAAAAAGCAATGGTCTTGAGTCAGACATGGGAAAAACGATGAAAATGGGACTTTTTA
TATTCCTTGTTTTACCCTCAACGTATCTTGACAACTTTGCTTCCAGATATTTCCCTCGGAGTGATGGGGAATACCTT
GTTGCATGGCACTGTGTGTTAACACATTACTTGCTCAACGGCCCCGTGACTTAATATCTACCGGAATGATGTTACAG
TTTTTATGGTATTCATTTGTCACACAATAATTTGTCCATGGCACTCCATCCACTGACTATTCGCAAGGTTGTGTGCC
TATAGCACTGCTTTGTGGGAATGTGTCACTCAGCGTGGATTAACTAACAGCAAAAAGTAATCGCCTCATTGTAATGT

ACAATTTCTGCTGAATTGCTGTTCAGTGGCAATGTAATAATATCTATGTAATCTTAATTTTGTAGCAATCTTGCAAG
AAAACAAAAATGACAAAAAAAGTTTGATAGCTAGAGGCCTGCCATGTACAGCATGTCATATGGTTCTATTTTTTTCT
CTGATGAAAATTAAAAAAAAAAGCAGCAAAATAAAAAAAAAAAAAAAAAAAAAAAAAATTATAAAAATGGGGGGGGGG
GGGGGGGG

>SWS2_TRINITY_DN3614_c0_g1_i3

AAGGAGGGAGCCTTCATCTGCTGTTCAACGAAGAGAGAATTTACACCGATACGGGTTGGACTCTTTTGCGATGCTGC
AGTAATCTACAGAGGGAGATAACACAATTGATTGAAGCTTGAAGGTGTTACAGCGTTCCTCGTGGAGTGGTTGCCAG
TTTCAAGCAAGATGAAGGCAGTACCCGAGTTTCACGATGACTTCTGGATCCCCATCCCTTTAGATACCAACAATATC
TCTGCTTACAGTCCTTTCCTGGTCCCCCAGGACCACCTGGGTCACCATGGCGTGTTCATGGCCATGGCTGCCCTTAT
GTTCTTCCTTTTCGTGGCAGGGACTGCGATCAACATCCTTACCATTGTTTGCACAATTCAATACAAGAAACTCAGAT
CTCACCTTAACTATATTCTTGTGAACCTTGCCGTTTCAAACCTGTGGGTGTCCGTTTTTGGTTCCTCGGTAGCATTC
TACGCCTTCTGGAGTAAATACTTTGTCTTCGGGGTGATAGGATGTAAAATTGAGGGCTTCACTTCAACAATTGGAGG
AATGGTGAGTTTGTGGTCTCTTGCTGTGGTGGCATTTGAAAGGTGGCTGGTCATTTGCAAACCCCTTGGGAACTTTA
CCTTCAAGACCACTCATGCCATAATTGGCTGCATACTTCCTTGGTGTATTGCATTGTCAGCTGGACTCCCTCCACTG
TTTGGATGGAGCCGGTACATCCCTGAAGGTTTGCAGTGCTCTTGTGGACCTGACTGGTATACGACTAACAACAAATA
CAACAACGAATCCTATGTCATGTTTTTGTTCTGCTTCTGCTTTGCGGTTCCTTTCACCACCATCGTGTTCTGTTATG
GTCAGCTGCTCATMACACTCAAATTRGCAGCCAAAGCTCAAGCAGATTCAGCTTCGACCCAGAAGGCAGAGAAGGAG
GTGACAAAGATGGTGGTGGTGATGGTGTTCGGCTTCTTGATATGCTGGGGACCATATGCTTGCTTTGCTCTCTGGGT
CATTTCCCACCGTGGTGAAACATTTGACCTGAGATTGGCAACCATACCATCCTGCCTTTGTAAAGCCTCAACAGTCT
ACAATCCTGTCATCTACGTCTTAATGAACAAACAGTTCCGTTCCTGTATGATGAAGATGGTCTGTGGCAAGAATATT
GAGGACGACGAGGCTTCTACTTCATCTCAGGTCACCCAGGTCTCCTCTGTTGCACCAGAGAAATAAACCCATTCTTA
ATGAAACCTCATCCTGCTGACAGAATCACTCTGCTGCATGGCAGTGAAAAATAATTTTTCAAAGATTGTAAATAATA
AACATAGACCCACTAGGAATTTTTGCTTTTAATGTCAACATTTTTGTAACTTGGTCTTGGCTTGCTGAACAAGTAGA
ATGAGTAGTTCATCTTTATCATTAAAAATGTACAATAAATGCAAAGAAAGAAAGAAACAATTGTGGTGTCGAATTTA
TCAGTTCAACAATTGTGACATACACGTTTAAATGAATTATATTACATTAGTCTCATTTCTTCCTGTTTTAAATGGAT
TGTTGTCAGCACACCATTTGACAACAACTAAAACATGAAAACAGAGAAGTATCTCGTATATCAGATTGTAATCAAGT
CCGTAGCTCTTTAGTCATGAGTAATGGAGTAACACATCCAGGGATGGTCTTGTACTTGATCCAATAAGTAGCAAGAT
CAAGATAAAAGTGCAAAGAAAGAATATGACATTAGCTTTTGTCATTTGACACAATAAATACAAGAACAATATTAGCA
CAGTTAATCAAGAAATTAAACCACAAATGATGTTAGAGAAACATTGAGCAGTGAAAGCAAAGTTTTGCCCATGTTAG
ATTAACATCTATTATATGGATTAATTCCATTCAAAAATGTTAAATAAATGTTCAGTACTTAATACCACGAATAAAGC
AGTTGTCCAATGGCAGACCGTGAGACCAAACTATTATCTTGAGGATTCGTTGAGAATTCAGTTGTATATCCGAGTTC
TGCTAGTAGTGAACTTTTGCCAATCACGAATAGTTCAGAACTAGCGGAACTCCAGACATAGGACTGACTGAGTGATG
CCCCTTGATACTTTCTGATCCATCTGTTTCTGAAAAACATCCTTTTCAACAACATCTCTAAATATATCTATGATTTA
CACTGTCATATAGCAACAAAAAAAACACTATAGTGACCCTTGACTCTCAATGCTGCACATTAATTCCAATAGTCTAC
TCAACTTCAATTTATTTAGATTAGCTTCCACTTTAAAACAAAGTGTGAAAGATCAAAACGAACAAATACTCGTTTTG
GATAAACACATACATCAGTACAATTCTCTGCAATTTTAAGATTGAAAAGGAAACAACTGATAGAGTAAAAAAAGTGT
ACAGACTGCTCTCGTCTTCGTGTTGGCTCCATCCTCACCCTGGTTTGGTCTGTTCACTTCCTCCTGCTCCACTTTGC
CACAGTAGATAGAGAAGCCGATGGCTGAAAAACAGAAAAAAAAAAAAAAAAAACAAAAAAGAATAATAAAAAAAAAAAT
TAAAAAAAAAAAACAAAAACCCCCAAAAAAATTAAAATAT

>CYP27C1_TRINITY_DN18543_c0_g1_i2

ATGGCTCTTCAAGCTACTATTCTACACATGGCCCGGACGAATCTGCTCCAGGAGTCATGCAAGCAGCTTCTCATCCA
AGCCCGTGGGCTGCACAAGTCCACGGCGAGCGGCTCTCTGGAGATCGCGGCGCACAGCCAGGCGGAACTGAAGGAAG
AAAACGAGGTGAGTCCGGCTGTGGTGGGGCTGAAGGAGACCACAGTGAAGACCCTCAAGGAGATGCCCGGACCCAGC
ACTATCTCGAACCTCGTTGAGTTCTTCTACCGGGATGGATTCAGCCGCATCCATGAGATCCAGACGGAGCACGCGCA
GAAGTATGGGAAGATCTTTAAATCTCGATTCGGACCTCAGCTGGTGGTGTCCATCGCGGACAGAGACTTGGTGGCAC
ATGTTCTCAGGTCTGAAGGTACGACTCCTCAGAGAGGCAACATGGAGTCCTGGAAGGAGTACAGAGATTTGAGGGGA
AGATCCACTGGACTCATCTCAGCCGAGGGTGATGAGTGGCTGAAGATGCGCAGCGTTCTGCGACAGCTGATTATGCG

GCCCAAAGATGTGTTTGCTTTCGCTCCTGATGTTAACGATGTGGTGGTCGACCTGGTGAAGAGAGTGAAGACGCTGC
GCAGCCAGCAGGACGATGGTCAGACTGTCCTTAACATCAATGACCTGTTCTTCAACTATGCCATGGAAGGTGTTGCA
ACCATTTTGTACGAGACTCGCCTGGGCTGCTTGGAGAATGAAATTCCCAAGATGAGTCAAGAGTACATCGCTGCGCT
GCATCTCATGTTCAGCTCTTTCAAGACAACCATGTATGCCGGCGCCATTCCCAAATGGCTGCGTCCCATAATTCCCA
AACCCTGGGAGGAATTCTGCAGCTCATGGGACGGACTCTTTAAATTCAGCCAGATCCATGTGGACAAGAAGCTTTCA
GAGATTAAAAAGAAGATGACTAAAGGTGAAGAAATTAAAGGAGGGTTGCTGACTCACATGCTGGTCACCAGAGAGAT
GAATCTAGAGGAGATCTACGCAAACATGACTGAAATGCTTCTGGCTGGAGTGGACACGACCTCTTTCACACTGTCAT
GGAGCACATACCTTCTGGCAAGACATCCCACAGTGCAGCAGGAGATTTTCGAGGAAGTGGACAGAGTGCTGGGCGGG
CGCGTCCCAACTGGAGAGGATGTTGCTTATCTGCCCCTTATTAGAGGGCTTGTCAAAGAGACGCTCAGGCTTTTTCC
CGTTCTCCCTGGAAATGGACGCGTTACACAGGATGATCTGATCGTCGGAGGTTATTTCATCCCTAAAGGGACTCAGC
TGGCTCTGTGTCATTACTCCACCTCAGTGGATGAGGAGAACTTTCCCCGTCCTGGAGAATTCCGTCCAGATCGCTGG
ATCCGCAAGGATGCCTCCGACCGCGTCGACAACTTCGGCTCCATCCCGTTTGGCTACGGCATCCGCAGTTGCATCGG
GAGGAGAATAGCTGAGCTCGAGATGCATCTGGCTCTCACACAGCTTCTACAGAAGTTCCACATTGAGGTGTCTTCCC
AGACCACCGATGTGCGTGCCAAGACTCATGGCCTGCTCTGTCCAGGTGCACCCATCAACCTTAGATTTGTGGACCGA
AAGTAATCTGTCATTACACTATGTATTCATATAAACCATAATCCACACTGTGTTTCTTGAAGTCTAAAGATGTCATT
GTGTATAATAGTGCCTCTTCTTGCTTTTATCTAACAAACAATGTTTTTTTTTATTTGTGTAAAGCGCCTCATTCAAA
GCAAGAACATATTTATTAGATAATTGTTAGTTTCTTGTTTGAATATTTATAGCCAATCAGCTGACCAAATATTTTTA
AAATGTGTAAACAGACACTATTTTATCGAGTCGGATCAGGACGAATTACTTTGAGAATTGTGTTAGAAAATTCTCGT
TTTCTTCACATGCATTTACATCAAACTAGCTGTGTAACTTTTGTCTCTTAGTTTTTTTTTTTGTTTCACTCCGCCTT
AAAATCATTGTGAAATATGGTCAAAAAGACCTTGTATATAGTTGTTTTACTATTAAGTTAGCTTTAGTGCTAAGCTC
ATTATTTATTTAACATTGTACGCTTATATACAGTCAAAGACTTCTAGCTTACAAGCAGGAAATTCCTAAAGCAAACA
TGAGTGAGTAAATCCTATTACAACTACAAAGCTACTTTAGAAGACCATGCTAATATCCTGTCCTGTGGACACTAATA
TTAGATATTGTTTAAAGGAAACACATCTAATCACAACAAAGCCCATCTTTATTTCAGTGTGGCTATCTCATTTGATA
TAACAGTGATGATTTTGTGTTGGTTCCACTAAAACACCAAAGGCAAAATTGTGTGCACTAGCCTTATAGGCAATGAA
TGGATAATGGGACATGCTAAAAAATGAGCTGCTTTTGGTTAACATGAATGTTGGTTTTGAGGGTTGTGAAGTAAGTA
CTACATTGTGTGATATTTGGAAGTTTTTGATATGTGGAAACAGGCTTTTGAATAAGTTACTGAAATTTGTATGAAGT
AAGTACTACATTGTGTG

>RHO_TRINITY_DN449_c0_g1_i1

GGAAGAGAGGGTAGCACGGTCCTGCCTCGTTTCTCTACATTCCTGCCAAGTCCTCCAAAGACCACCGCAGAAGGGGC
TGAGCACAACATCCAACCGCAGCCATGAACGGTACAGAGGGACCGGCATTCTACGTGCCTATGTCCAATGCCACCGG
CATTGTTAGGAGCCCCTACGAATATCCCCAGTACTACCTGGTGGCACCATGGGCGTACGCCTGCCTGGCCGCTTACA
TGTTCTTCCTCATCCTCACCGGCTTCCCCATCAATTTCCTCACTCTGTACGTCACCCTCGAGCACAAGAAGCTGCGC
ACGCCCCTCAACTACATCCTGCTGAACCTGGCCGTCGCCGATCTCTTCATGGTGTTCGGCGGCTTCACCACGACGAT
CTACACCTCCATGCACGGCTACTTCGTGCTGGGGCGCCTCGGCTGCAACATCGAAGGCTTCTTCGCSACCCTGGGYG
GTGAAATCGCGCTGTGGTCCATCGTCACGCTGGCCATCGAGAGGTGGCTGGTCGTCTGCAAGCCCATCACCAACTTC
CGCTTCGGAGAGGACCACGCCATCATGGGAGTGGTCTTCACGTGGGTCATGGCCAGCTCCTGCGCCGTGCCTCCCCT
GGTCGGCTGGTCCCGCTACATCCCCGAGGGCATGCAGTGCTCCTGCGGAGTCGACTACTACACCCGCGCCGAGGGCT
ACAACAACGAGTCCTTCGTCATCTACATGTTCCTTGTCCACGCCCTCATCCCGTTCTGCGTCATCTTCTTCTGCTAC
GGCCGCCTGGTCTGCACCGTGAAGGAAGCCGCTGCCCAGCAGCAGGAGTCCGAGACCACGCAGAGGGCCGAGCGCGA
GGTCACCCGCATGGTCATCCTCATGGGCTTCGCCTACCTGGTGTGTTGGTTGCCCTACGCCAGCGTGGCCTGGTACA
TCTTCACCCACAAGGGAACGGAATTCGGGCCGGTCTTCATGACAATCCCAGCCTTCTTTGCCAAGACCTCCGCCGTC
TACAACCCGCTCATCTACATCTGCATGAACAAGCAGTTCCGCCACTGCATGATCACCACCCTGTGCTGCGGCAAGAA
CCCCTTCGAGGAGGAAGAGGGCGCCTCCACCACCGCATCCAAGACCGAGGCTTCCTCCGTGTCGTCCGTGTCCCCGG
CATAAACACGCGGGCGAGACACACCTCGAGCAGCGACACGCACTGGGCTTCGACGGGCTTCAACCCACTCAGAGACC
ACGGAGCGCTCAGCCCAGGGAAACGAGCGACCGCTACCACTTGCAGAAAAAATCCTCTGTCAGTTTTCCTTTTTGTA
TTTTTACAAAACCCAATTGGTCCAACCAAAAGACAGTTCTGAGAGAGGACAGACCATGTCCCAGTTTCAGTACATCC
AGCGAGTCCAGCGTAATGGTACATACGATTCTTTTTGTTTTTCTTCCTAAAATGCAGCAAAAAGAAAAAATATCTTA
ACTCTTACGGCTGGACTCCTTATACTGGCTTTGTTGTGATTGTAGAGGCATGTATTCAAGGCAACGTAACAATAAAA

AGCACTTTGCAAATGAAAAAAAAAAAAAAAAAAAAAAAAAAAAAAAAAAAAAAAAAAAAAAAAAAAAAAAAAAAACC
CCAAAAAAGAAAAAACCAACAAATAAAAACCATAACCCAAAACCACAAAAAATTTTAACAAATAAAAATGAAAAA

Table S2. RNA sequencing samples, read depth, and percentage aligned.

| Sample ID | Site | Season | Sex | Total reads | Percent Aligned |
|---|---|---|---|---|---|
| LCV_1_24Jul20 | Lightning creek | Summer | Male | 21839574 | 64.3 |
| LCV_4_24Jul20 | Lightning creek | Summer | Male | 21958990 | 72.6 |
| LCV_7_24Jul20 | Lightning creek | Summer | Female | 22023320 | 72.7 |
| LCV_12_24Jul20 | Lightning creek | Summer | Female | 99146221 | 63.3 |
| LCV_2_21Nov20 | Lightning creek | Late Fall/Winter | Not determined | 28707701 | 56.7 |
| LCV_5_21Nov20 | Lightning creek | Late Fall/Winter | Male | 16365342 | 20.9 |
| LCV_9_21Nov20 | Lightning creek | Late Fall/Winter | Male | 16478292 | 50.6 |
| LCV_12_21Nov20 | Lightning creek | Late Fall/Winter | Male | 22983751 | 58.4 |
| PCJ_2_4Aug20 | Polecat creek | Summer | Male | 10735757 | 48.1 |
| PCJ_5_4Aug20 | Polecat creek | Summer | Female | 14375041 | 49.6 |

| | | | | | |
|---|---|---|---|---|---|
| PCJ_7_4Aug20 | Polecat creek | Summer | Female | 16011929 | 26.8 |
| PCJ_10_4Aug20 | Polecat creek | Summer | Male | 14001305 | 37.8 |
| PCJ_4_20Dec20 | Polecat creek | Late Fall/Winter | Not determined | 19651629 | 49.6 |
| PCJ_7_20Dec20 | Polecat creek | Late Fall/Winter | Not determined | 19663780 | 49.7 |
| PCJ_8_20Dec20 | PCJ | Late Fall/Winter | Not determined | 18991969 | 50.2 |
| PCJ_9_20Dec20 | PCJ | Late Fall/Winter | Male | 22898984 | 64.4 |

Figure S3. Representative (a) HPLC chromatograms and (b-c) UV-Vis spectra of selected peaks ocular retinoid extracts of *C. lutrensis* sampled from two Oklahoma creeks.

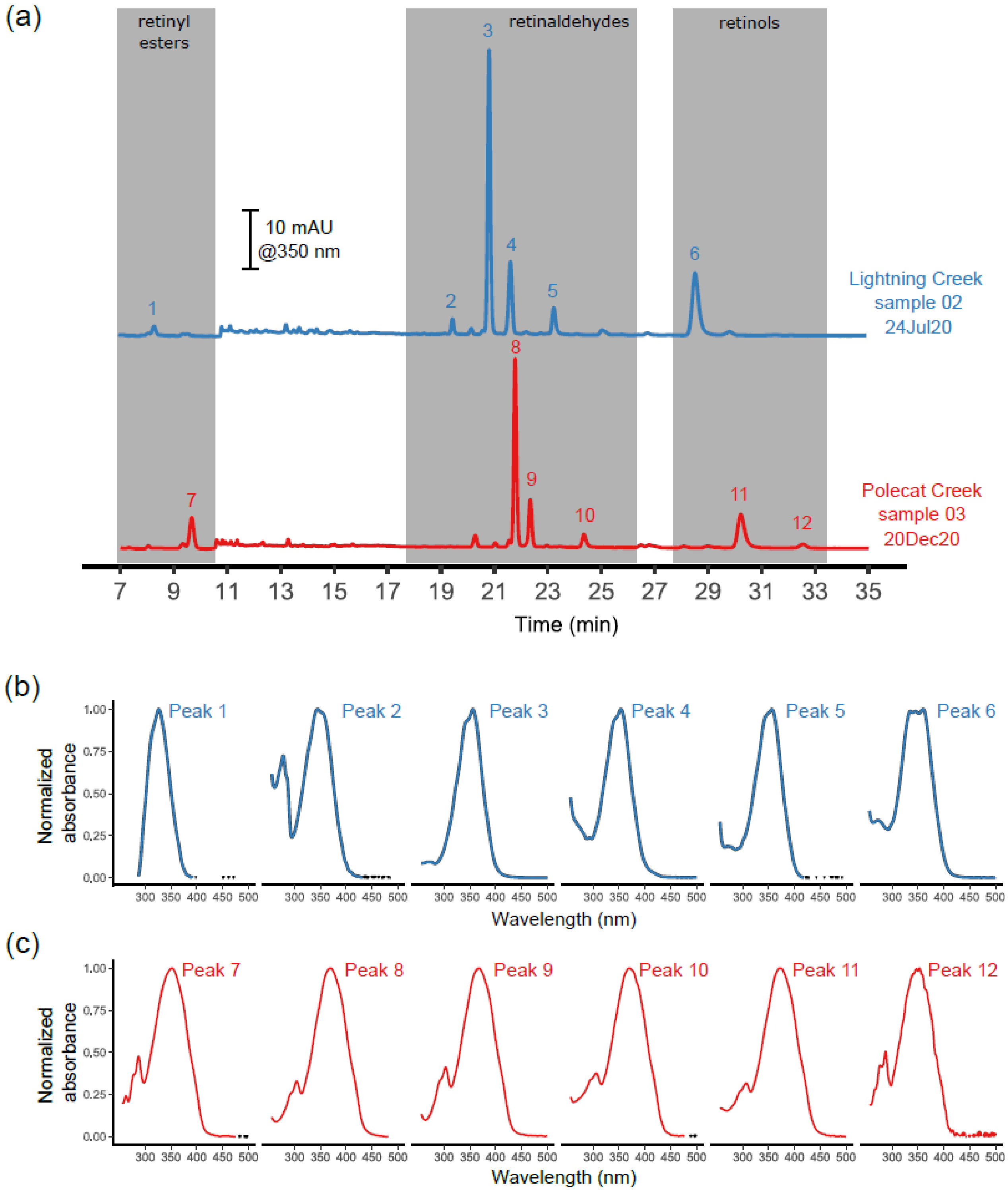

Figure S4. A pairwise alignment of the amino acid sequence of *Danio rerio* and *Cyprinella lutrensis* cytochrome P450 27c1(CYP27C1).

```
Danio_rerio             MALQSTILHMARKNLLQESCRQLLIQTHGLHKSVASGSLEIAAHSQADLKEESAVSPAEE    60
Cyprinella_lutrensis    MALQATILHMARTNLLQESCKQLLIQARGLHKSTASGSLEIAAHSQAELKEENEVSPAVV    60
                        ****:*******.*******:*****::*****.*************:****. ****

Danio_rerio             VQKAARVKSLKEMPGPSTVANLLEFFYRDGFSRIHEIQMEHAKKYGKIFKSRFGPQFVVS    120
Cyprinella_lutrensis    GLKETTVKTLKEMPGPSTISNLVEFFYRDGFSRIHEIQTEHAQKYGKIFKSRFGPQLVVS    120
                          * : **:*********::**:*************** ***:*************:***

Danio_rerio             IADRDMVAQVLRSESATPQRGNMESWKEYRDLRGRSTGLISAEGDEWLKMRSVLRQLIMR    180
Cyprinella_lutrensis    IADRDLVAHVLRSEGTTPQRGNMESWKEYRDLRGRSTGLISAEGDEWLKMRSVLRQLIMR    180
                        *****:**:*****.:********************************************

Danio_rerio             PRDVAVFSSDVNDVVADLVKRVKTLRSQQDDSQTVLNINDLFFKYAMEGVATILYETRLG    240
Cyprinella_lutrensis    PKDVFAFAPDVNDVVVDLVKRVKTLRSQQDDGQTVLNINDLFFNYAMEGVATILYETRLG    240
                        *:** .*: ******.***************.***********:****************

Danio_rerio             CLENEIPKMSQEYITALHLMFSSFKTTMYAGAIPKWLRPIIPKPWEEFCSSWDGLFKFSQ    300
Cyprinella_lutrensis    CLENEIPKMSQEYIAALHLMFSSFKTTMYAGAIPKWLRPIIPKPWEEFCSSWDGLFKFSQ    300
                        *************:**********************************************

Danio_rerio             IHVDKRLSEIKKQMEKSEEIKGGLLTHMLVTREMNLEEIYANMTEMLLAGVDTTSFTLSW    360
Cyprinella_lutrensis    IHVDKKLSEIKKKMTKGEEIKGGLLTHMLVTREMNLEEIYANMTEMLLAGVDTTSFTLSW    360
                        *****:******:* *.*******************************************

Danio_rerio             STYLLARHPTIQQQIFEEVDRVLGGRVPTGEDVPYLPLIRGLVKETLRLFPVLPGNGRVT    420
Cyprinella_lutrensis    STYLLARHPTVQQEIFEEVDRVLGGRVPTGEDVAYLPLIRGLVKETLRLFPVLPGNGRVT    420
                        **********:**:******************** *************************

Danio_rerio             HDDLIVGGYLIPKGTQLALCHYSTSMDEENFPRPEEFRPDRWIRKDASDRVDNFGSIPFG    480
Cyprinella_lutrensis    QDDLIVGGYFIPKGTQLALCHYSTSVDEENFPRPGEFRPDRWIRKDASDRVDNFGSIPFG    480
                        :********:***************:******** *************************

Danio_rerio             YGIRSCIGRRIAELEMHLALTQLLQNFHIEVSPQTTEVHAKTHGLLCPGASINLRFTDRK    540
Cyprinella_lutrensis    YGIRSCIGRRIAELEMHLALTQLLQKFHIEVSSQTTDVRAKTHGLLCPGAPINLRFTDRK    540
                        *************************:****** ***:*:*********** *********
```

Table S3. Results of AICc analysis (Mazerolle 2020) of models using site as a factor or our direct measurements of light environment to predict the proportion of $A_2$ chromophore in the eyes of red shiners sampled at three sites throughout the year.

| Model | K | AICc | Delta AICc |
|---|---|---|---|
| `A2 proportion ~ site*season + eye diameter` | 14 | -420.0 | 0.00 |
| `A2 proportion ~ red:blue ratio*season + eye diameter` | 4 | -317.6 | 102.4 |
| `A2 proportion ~ irradiance*season + eye diameter` | 4 | -238.2 | 181.8 |

Mazerolle, M. J. 2020. "Model Selection and Multimodel Inference Using the AICcmodavg Package." *R Vignette*. https://mirrors.cloud.tencent.com/CRAN/web/packages/AICcmodavg/vignettes/AICcmodavg.pdf.
